# Comparative transcriptomics analyses across species, organs and developmental stages reveal functionally constrained lncRNAs

**DOI:** 10.1101/607200

**Authors:** Fabrice Darbellay, Anamaria Necsulea

## Abstract

**Background:** Transcription of long non-coding RNAs (lncRNAs) is pervasive, but their functionality is disputed. As a class, lncRNAs show little selective constraint and negligible phenotypic effects upon perturbation. However, key biological roles were demonstrated for individual lncRNAs. Most validated lncRNAs were implicated in gene expression regulation, in pathways related to cellular pluripotency, differentiation and organ morphogenesis, suggesting that functional lncRNAs may be more abundant in embryonic development, rather than in adult organs.

**Results:** Here, we perform a multi-dimensional comparative transcriptomics analysis, across five developmental time-points (two embryonic stages, newborn, adult and aged individuals), four organs (brain, kidney, liver and testes) and three species (mouse, rat and chicken). Overwhelmingly, lncRNAs are preferentially expressed in adult and aged testes, consistent with the presence of permissive transcription during spermatogenesis. LncRNAs are often differentially expressed among developmental stages and are less abundant in embryos and newborns compared to adult individuals, in agreement with a requirement for tighter expression control and less tolerance for noisy transcription early in development. However, lncRNAs expressed during embryonic development show increased levels of evolutionary conservation, both in terms of primary sequence and of expression patterns, and in particular at their promoter regions. We find that species-specific lncRNA transcription is frequent for enhancer-associated loci and occurs in parallel with expression pattern changes for neighboring protein-coding genes.

**Conclusions:** We show that functionally constrained lncRNA loci are enriched in developing organ transcriptomes, and propose that many of these loci may function in an RNA-independent manner.

## Background

Long non-coding RNAs (lncRNAs, loosely defined as transcripts that lack protein-coding potential, at least 200 nucleotides long) are an excellent illustration of the ongoing conceptual tug-of-war between biochemical activity and biological function (Graur et al. 2013; Doolittle 2018). The development of sensitive transcriptome exploration techniques led to the identification of thousands of lncRNA loci in vertebrates (Guttman et al. 2009; Khalil et al. 2009; Iyer et al. 2015; Pertea et al. 2018). While this ever wider class of transcripts includes well-studied lncRNAs with undisputed biological roles, such as *Xist* (Brown et al. 1991) or *H19* (Brannan et al. 1990), experimental validations are lacking for the great majority of lncRNAs and their functionality is controversial.

The first functional characterizations of individual lncRNAs forged the idea that these non-coding transcripts are important contributors to gene expression regulatory networks. This has been unequivocally proven for some lncRNAs, such as *Xist*, whose transcription and subsequent coating of the X chromosome triggers a complex chain of molecular events leading to X inactivation in placental mammals (Gendrel and Heard 2014). Additional proposed mechanisms for gene expression regulation by lncRNAs included directing chromatin-modifying complexes at specific genomic locations, to control gene expression in *trans* (Rinn et al. 2007); providing decoy targets for microRNAs (Cesana et al. 2011); enhancing expression of neighboring genes through an RNA-dependent mechanism (Ørom et al. 2010). These initial studies generally asserted that the biological function of lncRNA loci is directly carried out by the transcribed RNA molecule. However, it was shown early on that in some cases the function resides in the act of transcription at a given genomic location, rather than in the product of transcription (Latos et al. 2012). Moreover, several recent publications showed that lncRNA transcripts are not required, and that instead biological functions are carried out by other elements embedded in the lncRNA genomic loci (Bassett et al. 2014). For example, it was recently shown that transcription of the *Linc-p21* gene, originally described as a *cis*-acting enhancer lncRNA, is not needed to regulate neighboring gene expression (Groff et al. 2016). Genetic engineering of multiple lncRNA loci in mouse likewise showed that lncRNA transcripts are dispensable, and that gene expression regulation by lncRNA loci is instead achieved by the process of lncRNA transcription and splicing, or by additional regulatory elements found in lncRNA promoters (Engreitz et al. 2016; Anderson et al. 2016). Furthermore, some attempts to look for lncRNA function through genetic engineering approaches showed that the tested lncRNA loci are altogether dispensable (Amândio et al. 2016; Zakany et al. 2017; Goudarzi et al. 2019). These recent observations signal a paradigm shift in lncRNA biology, as it is increasingly acknowledged that, even when phenotypic effects can be unambiguously mapped to lncRNA loci, the underlying biological processes are not necessarily driven by the lncRNA transcripts themselves.

Importantly, this new perspective on lncRNA biology had been predicted by evolutionary analyses, which have long been used to evaluate the functionality of diverse genomic elements (Haerty and Ponting 2014; Ulitsky 2016). Evolutionary studies of lncRNAs in vertebrates all agree that the extent of selective constraint on lncRNA primary sequences is very low, though significantly above the genomic background (Ponjavic et al. 2007; Kutter et al. 2012; Necsulea et al. 2014; Washietl et al. 2014; Hezroni et al. 2015). These observations are compatible with the hypothesis that many of the lncRNAs detected with sensitive transcriptomics techniques may be non-functional noise (Ponjavic et al. 2007), but may also indicate that lncRNA functionality does not reside in the primary transcribed sequence. In contrast, mammalian lncRNA promoters show higher levels of sequence conservation, similar to protein-coding gene promoters (Necsulea et al. 2014), as expected if they carry out enhancer-like regulatory functions independently of the transcribed RNA molecule. Moreover, it was previously reported that, in multi-exonic lncRNAs, splicing regulatory elements are more conserved than the rest of the exonic sequence (Schüler et al. 2014; Haerty and Ponting 2015), which is compatible with the recent finding that lncRNA splicing can contribute to neighboring gene regulation (Engreitz et al. 2016). Thus, detailed evolutionary analyses of lncRNA loci can bring important insights into their functionality, and can help to prioritize candidates for experimental validation.

At present, comparative transcriptomics analyses in vertebrates indicate that the extent of evolutionary conservation of lncRNA sequences and expression patterns is very limited. However, these studies were so far restricted to adult organ transcriptomes. In particular, it was shown that most known vertebrate lncRNAs are active in adult testes and thus likely during spermatogenesis, a process characterized by a permissive chromatin environment, which can promote non-functional transcription (Soumillon et al. 2013). The resulting lncRNA datasets may thus be enriched in non-functional transcripts. Additional lines of evidence suggest that the search for functional lncRNAs should be extended beyond adult organ transcriptomes. For example, involvement in developmental phenotypes was proposed for many experimentally-tested lncRNAs (Sauvageau et al. 2013; Ulitsky et al. 2011; Grote et al. 2013), and an enrichment for developmental transcription factor binding was reported for the promoters of highly conserved lncRNAs (Necsulea et al. 2014). These observations motivated us to add a temporal dimension to comparative lncRNA transcriptomics studies. Therefore, here we characterize the lncRNA transcriptomes of two model mammalian species (mouse and rat), in four major organs (brain, kidney, liver and testes), across five developmental stages that cover the entire lifespan of the individuals (including two embryonic stages, newborn, young adult and aged individuals). To gain a deeper evolutionary perspective, we generate similar data for embryonic stages of chicken somatic organs. We analyze the spatial and temporal expression patterns of protein-coding and lncRNA genes, in conjunction with their evolutionary conservation. We find that, while lncRNAs are overall poorly conserved among species in terms of primary sequence or expression patterns, higher frequencies of evolutionarily constrained lncRNAs are observed in embryonic transcriptomes. For many of these loci, biological function may be RNA-independent, as the highest levels of sequence conservation are observed on promoter regions and on splice signals, rather than on lncRNA exonic sequence. Our results are thus compatible with unconventional, RNA-independent functions for lncRNA loci, in particular for those that are expressed during embryonic development.

## Results

### Comparative transcriptomics across species, organs and developmental stages

To study protein-coding and lncRNA expression patterns across both developmental and evolutionary time, we generated RNA-seq data for mouse and rat, for four major organs (brain, kidney, liver and testes) and five developmental time points, including two embryonic stages, newborn, young and aged adult individuals (Figure 1A, Supplementary Table 1, Methods). The selected time points allow us to obtain a broad view of major organ ontogenesis and to capture drastic physiological changes during development (Theiler 1989). We chose to include in our study both young adult (8-10 weeks old) and aged adult individuals (12 to 24 months old), to investigate transcriptomic changes that occur later in life, thus completing our overview of the temporal patterns of gene expression variation. At the earliest embryonic stage (day 13.5 post-conception for mouse, day 15 for rat), only three of the four studied organs, with the exception of the testes, are well differentiated and large enough to be readily dissected. Our experimental design for mouse and rat thus comprises 19 organ / developmental stage combinations. Although most of our study relies on mouse-rat comparisons, to obtain a broader evolutionary perspective we generated comparable RNA-seq data for the chicken, for the two earliest developmental stages (Figure 1A, Supplementary Table 1). We obtained between 2 and 4 biological replicates for each species/organ/developmental stage combination (Supplementary Table 1). Additional RNA-seq samples from previous publications were included in the lncRNA annotation pipeline, to increase detection sensitivity (Supplementary Table 2, Methods).

**Figure 1.**
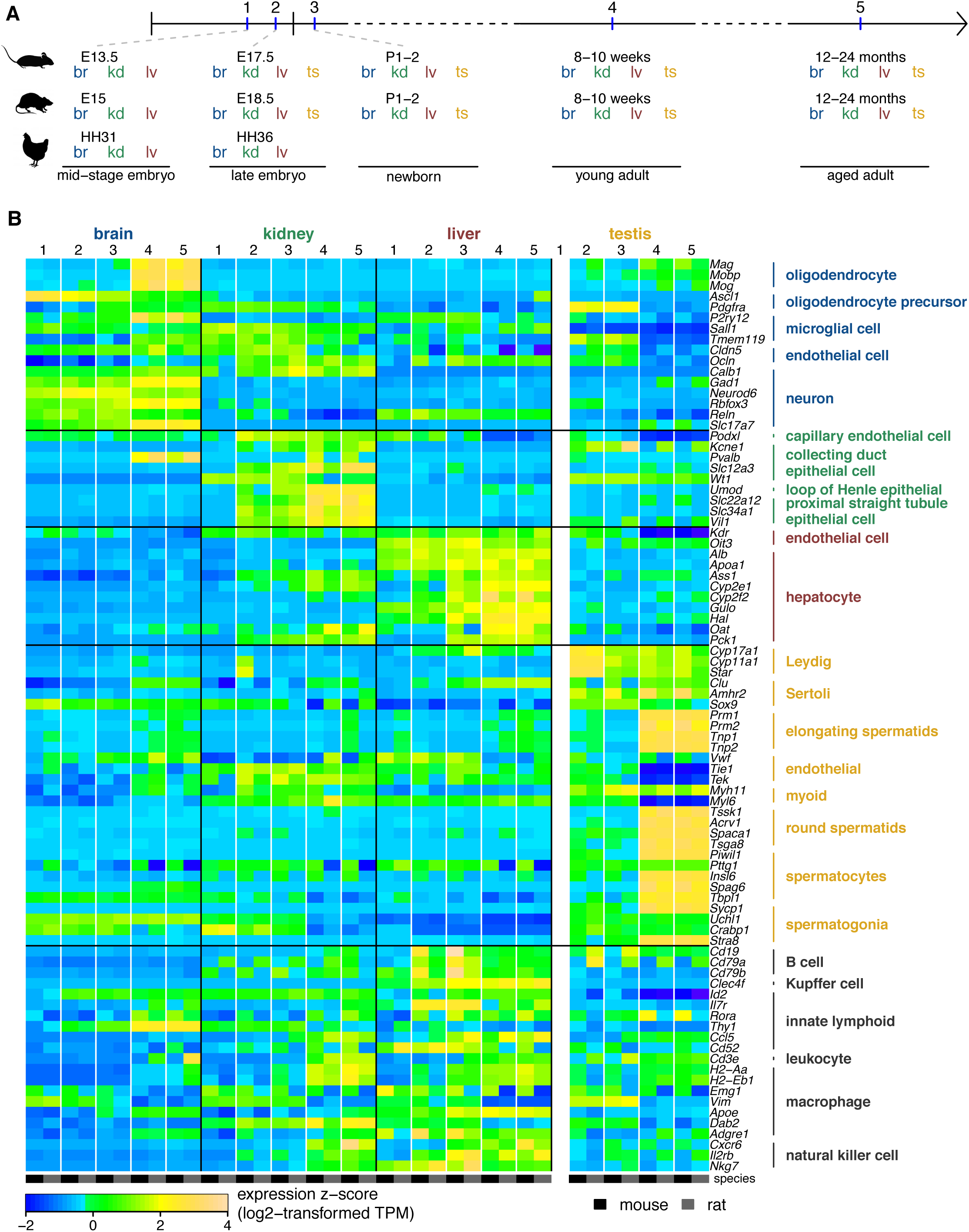
Comparative transcriptomics across species, organs and developmental stages. **A.** Experimental design. The developmental stages selected for mouse, rat and chicken are marked on a horizontal axis. Organs sampled for each species and developmental stage are shown below. Abbreviations: br, brain; kd, kidney; lv, liver; ts, testes. **B.** Expression of cell type-specific markers derived from single-cell experiments (full list provided in Supplementary Table 3), in our mouse and rat RNA-seq samples, averaged across biological replicates. The heatmap represents centered and scaled log2-transformed TPM levels (z-score). Developmental stages are indicated by numeric labels, 1 to 5. Species are color-coded, shown below the heatmap.

The organs and developmental stages included in our study differ greatly in terms of their cellular composition diversity. To verify that our whole-organ RNA-seq data reflects cellular composition heterogeneity, we assessed the expression patterns of cell population markers derived from single-cell transcriptomics studies (Tabula Muris Consortium 2018; Green et al. 2018) in our samples (Figure 1B, Supplementary Table 3). This analysis confirms that our transcriptome collection reflects expected developmental patterns. For example, mature oligodendrocyte cell markers are systematically highly expressed in adult brain, while oligodendrocyte precursor markers are more highly expressed in the earliest developmental stages (Figure 1B). Similarly, *Neurod6,* a gene involved in neuronal differentiation (Kathleen Baxter et al. 2009), is preferentially expressed in embryonic and newborn brain. Moreover, spermatogenesis-specific markers are enriched in adult but not in embryonic and newborn testes, while markers for somatic cells (Leydig, Sertoli cells) are expressed earlier during testes development (Figure 1B). Immune cell markers tend to be more broadly shared across organs and developmental stages, but show strongest expression in the late embryo and newborn liver (Figure 1B), consistent with this organ’s crucial role in establishing immunity (Nakagaki et al. 2018). In general, adult organ transcriptomes contain higher numbers of expressed cell type-specific markers (Figure 1B). However, as these genes were defined based on adult organ data, this observation may indicate that cell sub-populations that are specific to embryonic organs are under-represented in this marker set, rather than reflecting the true cellular diversity at different developmental stages.

We note that, in some cases, the cell type-specific markers predicted by single-cell transcriptomics studies have seemingly unexpected expression patterns in our whole-organ RNA-seq collection. For example, the expression of *Parvalbumin* (*Pvalb*), which was proposed as a marker for collecting duct epithelial cells in the kidney (Tabula Muris Consortium 2018), is highest in the adult and aged brain, for both mouse and rat (Figure 1B). Likewise, cellular retinoic acid binding protein 1 (*Crabp1*), which was predominantly detected in spermatogonia in a single-cell transcriptomics study of mouse testes (Green et al. 2018), is preferentially expressed in mid-stage embryonic kidney in our samples (Figure 1B). These apparent discrepancies likely reflect the pleiotropic nature of genes, as well as the presence of similar cell types across organs with distinct physiological functions (Arendt et al. 2016). In general, the genes proposed as markers for major cell types behave similarly in mouse and rat, although some species-specific patterns can be observed, in particular for immune cell markers (Figure 1B). Likewise, for those genes that had orthologues in the chicken, expression patterns are generally similar among species, with higher between-species divergence for immunity-related genes (Supplementary Figure 1). This observation confirms that the organs and developmental stages selected for our integrative transcriptomics study are comparable across species.

We note that the cell-type specific markers that we analyzed here cannot always discriminate between organs and developmental stages, as expected given that some cell sub-populations are shared among tissues. To further characterize our transcriptome collection and identify the patterns that are specific to each sample, we sought to identify genes that could serve as markers for organ/developmental stage combinations. To do this, we selected genes that have narrow expression distributions, and for which the maximum expression is observed in the same organ/developmental stage combination in mouse and rat (Methods, Supplementary Table 4, Supplementary Figures 2-5). Gene ontology enrichment analyses for the resulting lists of genes (Supplementary Dataset 3) are coherent with the cellular composition and biological processes at work in the corresponding samples. For example, we observe strong over-representation of genes involved in forebrain neuron differentiation in the mid-stage embryonic brains, while processes related to synaptic transmission are enriched among genes specifically expressed in adult brains (Supplementary Figure 2). In the kidney, the early developmental stages are expectedly enriched in genes involved in metanephric development (Supplementary Figure 3). The newborn liver stands out due to its strong enrichment in genes involved in immune response, while metabolic processes are over-represented in the adult livers (Supplementary Figure 4). Embryonic testes samples express genes implicated in gamete generation and gene silencing by miRNAs, including the *Piwi-*like genes, while adult testes transcriptomes are dominated by genes involved in spermatogenesis (Supplementary Figure 5). These expected patterns confirm that our whole-organ transcriptomics collection provides a good overview of the cell composition changes and physiological transitions that occur during organ development.

### Developmental expression patterns are well conserved among species for protein-coding genes

Broad patterns of transcriptome evolution are already visible in our analyses of cell type specific markers and of transcriptome complexity: individual gene expression profiles and numbers of expressed genes are generally similar between mouse and rat, while more divergence is observed between the two rodent species and the chicken (Figure 1B-D, Supplementary Figure 1). To further explore the evolution of developmental gene expression patterns, we performed a principal component analysis (PCA) on normalized, log-transformed TPM values for 10,363 protein-coding genes shared among the three species (Methods, Figure 2A). This analysis revealed that the main source of gene expression variability among species, organs and developmental stages is the distinction between adult and aged testes and the other samples, which are separated on the first PCA axis (Figure 2A). In contrast, embryonic and newborn testes are grouped with kidney samples from similar developmental stages, in agreement with the common developmental origin of the kidney and the gonads (McMahon 2016). The first axis of the PCA, which explains 67% of the total expression variance, also correlates with the developmental stage: samples derived from adult and aged individuals have higher coordinates on this axis than embryonic and newborn samples, for mouse and rat (Figure 2A). The second PCA axis (10% explained variability) mainly reflects the difference between brain and the other organs (Figure 2A). While mouse and rat samples are generally undistinguishable, the PCA confirms that there is considerably higher expression divergence between chicken and the two rodent species (Figure 2A). However, differences among major organs are stronger than differences among species, even at these broad evolutionary distances: brain samples all cluster together, irrespective of the species of origin, and are clearly separated from kidney and liver samples on the second PCA axis (Figure 2A). Interestingly, within the brain cluster, embryonic chicken samples tend to be closer to adult and aged rodent brains than to embryonic or neonate samples (Figure 2A).

**Figure 2.**
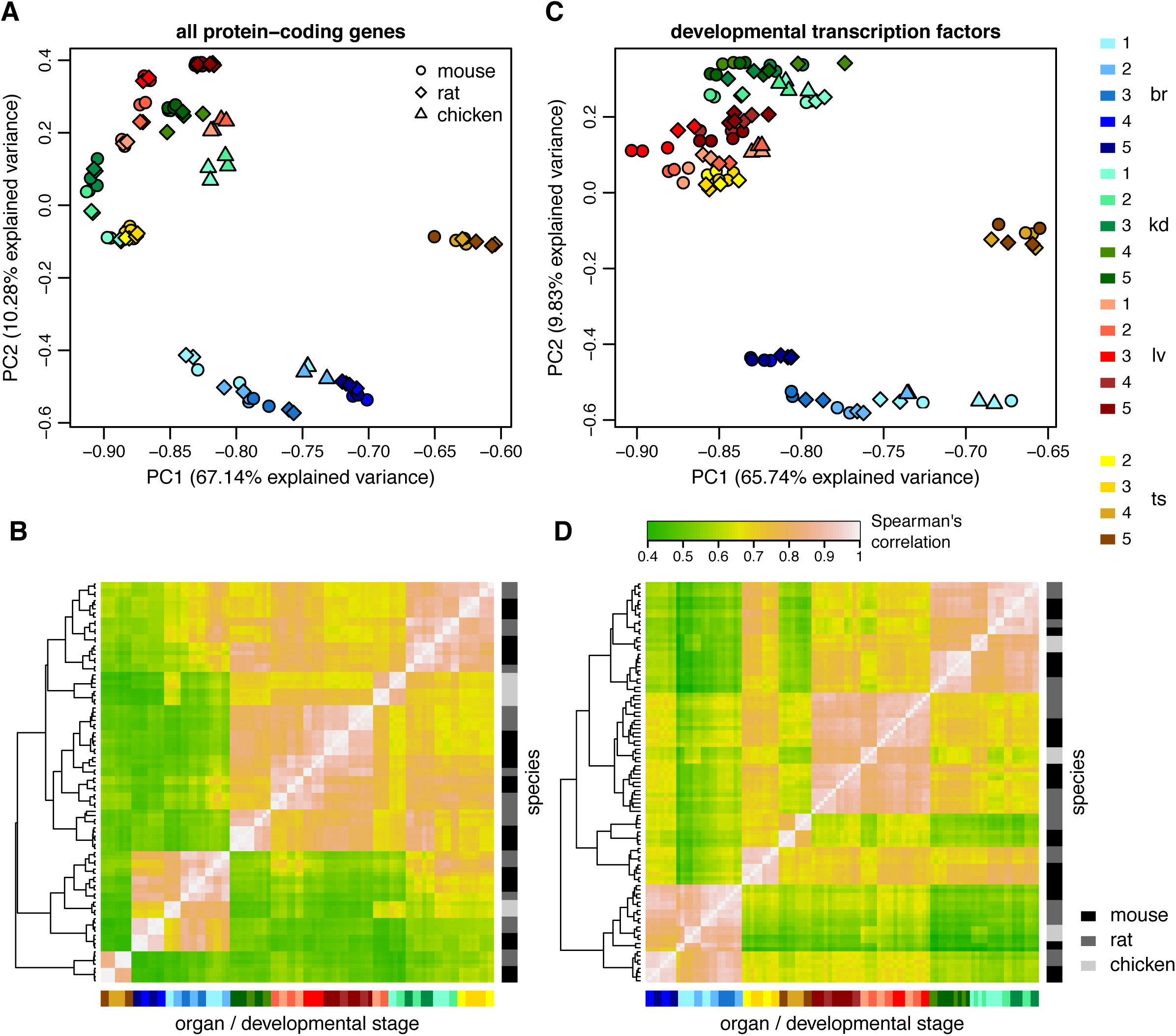
Protein-coding gene expression patterns are conserved across species. **A.** First factorial map of a principal component analysis, performed on log2-transformed TPM values, for 10,363 protein-coding genes with orthologues in mouse, rat and chicken. Colors represent different organs and developmental stages, point shapes represent different species. **B.** Hierarchical clustering, performed on a distance matrix derived from Spearman correlations between pairs of samples, for 10,363 protein-coding genes with orthologues in mouse, rat and chicken. Organ and developmental stages are color-coded, shown below the heatmap. Species of origin is color-coded, shown on the right. Sample clustering is shown on the left. **C.** Same as A, for 289 protein-coding genes with orthologues in mouse, rat and chicken, associated with organism development and regulation of gene expression in mouse gene ontology (Methods). **D.** Same as B, for 289 protein-coding genes associated with organism development and regulation of gene expression.

These broad patterns of gene expression variations among species, organs and developmental stages are confirmed by a hierarchical clustering analysis based on Spearman’s correlation coefficients between pairs of samples (Figure 2B). The strongest clustering is observed for adult and aged testes samples, followed by a robust grouping of brain samples, irrespective of the species (Figure 2B).

The grouping among samples derived from similar organs and developmental stages, irrespective of the species of origin, is considerably stronger for genes that are associated with embryonic development and with gene expression regulation (Methods, Figure 2C,D). For this set of genes, both the principal component analysis and the hierarchical clustering analysis are characterized by a near-perfect separation of organs and developmental stages, for all three species (Figure 2C,D). Chicken samples, which clustered apart from rodent samples in whole transcriptome analyses, are now grouped with the corresponding organs and developmental stages from mouse and rat. Our transcriptome collection can thus reveal highly conserved expression patterns for key regulators of embryonic development, across vertebrates.

### Variations in transcriptome complexity among organs and developmental stages

We next sought to assess the transcriptome complexity in different organs across developmental stages, for both protein-coding genes and lncRNAs. To predict lncRNAs, we used the RNA-seq data to reconstruct gene models with StringTie (Pertea et al. 2015), building on existing genomic annotations (Cunningham et al. 2019). We verified the protein-coding potential of newly annotated transcripts, based on the codon substitution frequency score (Lin et al. 2007, 2011) and on sequence similarity with known proteins, and we applied a stringent series of filters to reduce contaminations from un-annotated protein-coding UTRs and other artefacts (Methods). We thus obtain a total of 18,858 candidate lncRNAs in the mouse, 20,159 in the rat and 5,496 in the chicken, including both newly-annotated and previously known lncRNAs transcribed in our samples (Supplementary Dataset 1). We note that many of these candidate lncRNAs are expressed at very low levels. When imposing a minimum normalized expression level (transcript per million, or TPM) at least equal to 1, in at least one sample, the numbers of candidate lncRNAs falls to 12,199, 15,319 and 2,892 in the mouse, rat and chicken, respectively (Supplementary Datasets 2-3, Supplementary Table 4).

The differences in lncRNA content among species may reflect discrepancies in RNA-seq read coverage and sample distribution, as well as genome sequence and annotation quality. To correct for the effect of RNA-seq read coverage, we down-sampled the RNA-seq data to obtain the same number of uniquely mapped reads for each organ/developmental stage combination within each species (Methods). After this equalizing procedure, the number of detectable protein-coding genes (supported by at least 10 uniquely mapped reads) still shows broad variations among organs and developmental stages, with the highest numbers of genes detected in the testes, for all time points (Figure 3A). Large numbers of protein-coding genes (between 12,800 and 16,700) are detected in all samples. In contrast, for lncRNAs, the pattern is much more striking: the young and aged adult testes express between 11,000 and 12,000 lncRNAs, in both mouse and rat, while in somatic organs and earlier developmental stages we can detect only between 1,800 and 4,800 lncRNAs (Figure 3B). This observation is in agreement with previous findings indicating that the particular chromatin environment of the adult testes, and in particular of spermatogenesis-specific cell types, is extraordinarily permissive to transcription (Soumillon et al. 2013). Interestingly, the numbers of protein-coding genes detectable in each organ also varies among developmental stages. In young and aged adult individuals, the brain shows the second-highest number of expressed protein-coding genes, after the testes, as previously observed (Soumillon et al. 2013; Ramsköld et al. 2009). However, in embryonic and newborn samples, the kidney expresses higher numbers of protein-coding genes than the brain (Figure 3B).

**Figure 3.**
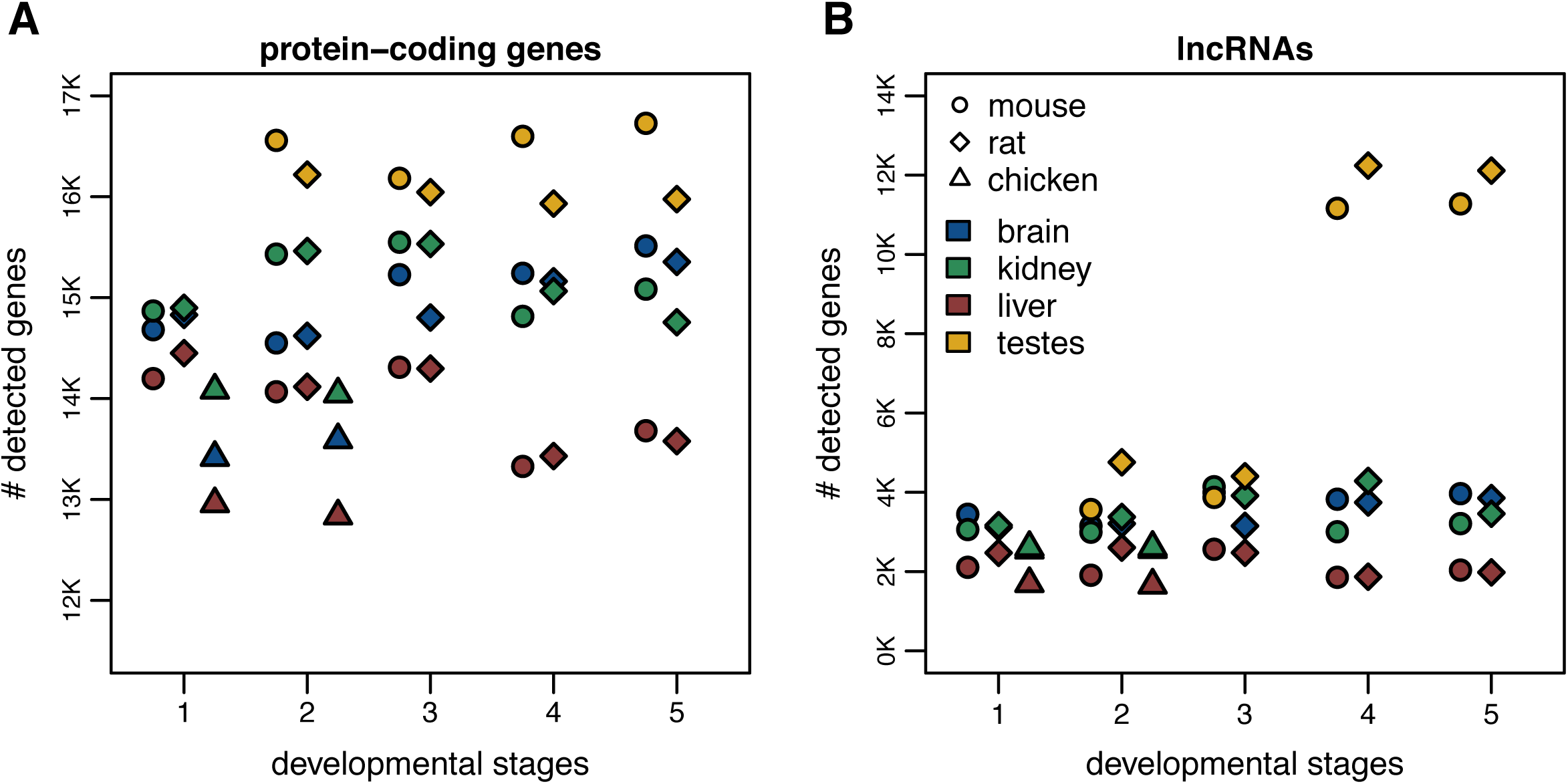
Transcriptome complexity in different species, organs and developmental stages. **A.** Number of protein-coding genes supported by at least 10 uniquely mapped reads in each sample, after read resampling to homogenize coverage (Methods). Colors represent different organs, point shapes represent different species. Developmental stages are indicated by numeric labels, 1 to 5, on the X-axis. We analyzed a total of 19,356 protein-coding genes in the mouse, 19,274 in the rat and 15,509 in the chicken. **D.** Number of lncRNAs supported by at least 10 uniquely mapped reads in each sample, after read resampling to homogenize coverage. We analyzed a total of 18,858 lncRNAs in the mouse, 20,159 in the rat and 5,496 in the chicken.

### Spatial and temporal expression pattern differences between protein-coding genes and lncRNAs

We next compared spatial and temporal expression patterns between protein-coding genes and lncRNAs. In agreement with previous findings (Soumillon et al. 2013), we show that lncRNAs are overwhelmingly preferentially expressed in the testes (Figure 4A). Indeed, more than 68% of lncRNAs reach their maximum expression level in this organ, compared to only approximately 32% of protein-coding genes, for both mouse and rat (Figure 4A). Interestingly, more than 80% of lncRNAs are preferentially expressed in young and aged adult samples, compared to only 62% of protein-coding genes (Figure 4B).

**Figure 4.**
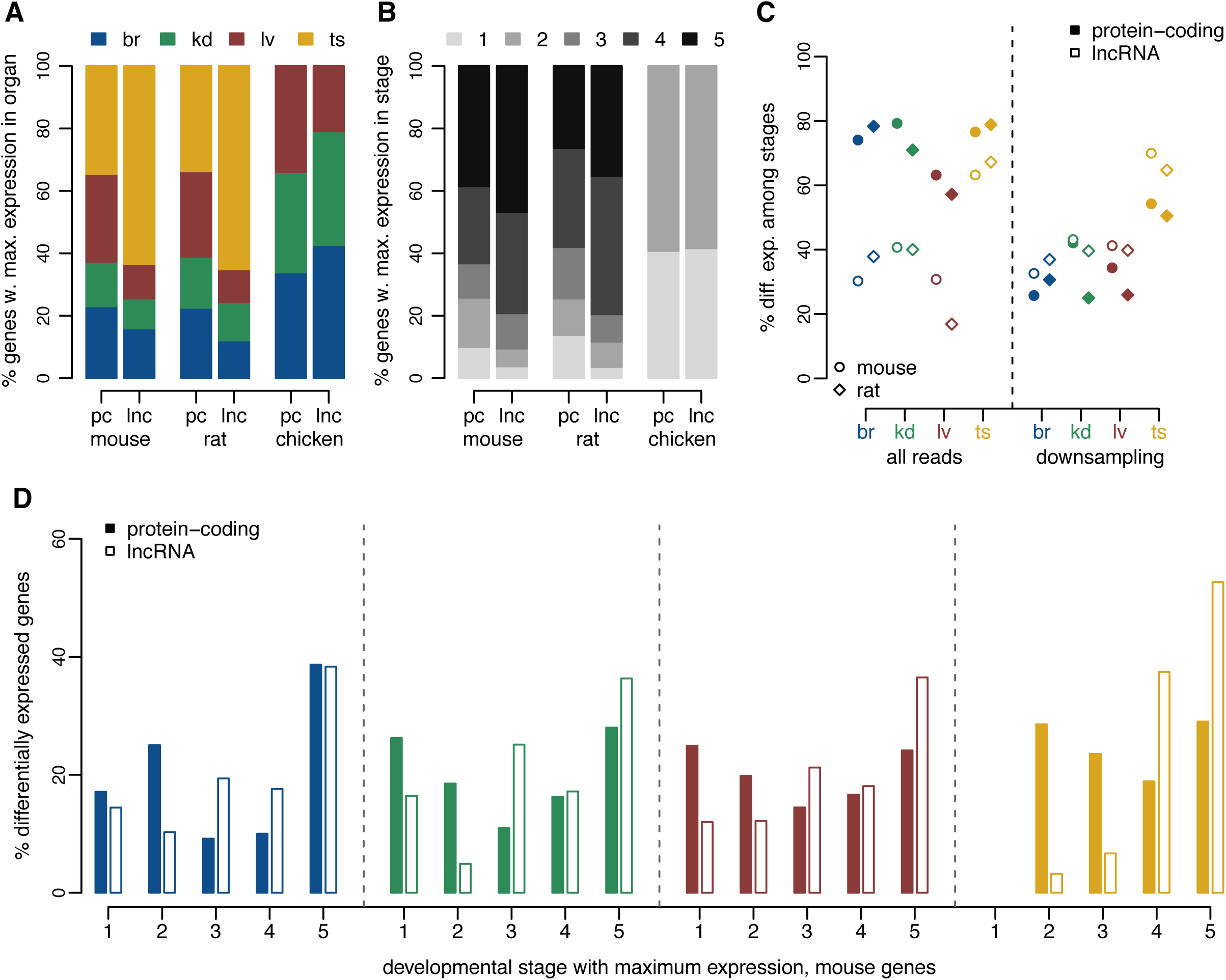
Different expression patterns for protein-coding genes and lncRNAs. **A.** Distribution of the organ in which maximum expression is observed, for protein-coding genes (pc) and lncRNAs (lnc), for mouse, rat and chicken. Organs are color-coded, shown above the plot. **B.** Distribution of the developmental stage in which maximum expression is observed, for protein-coding genes and lncRNAs, for mouse, rat and chicken. Developmental stages are color-coded, shown above the plot. **C.** Percentage of protein-coding and lncRNA genes that are significantly (FDR<0.01) DE among developmental stages, with respect to the total number of genes tested for each organ. Left panel: differential expression analysis performed with all RNA-seq reads. Right panel: differential expression analysis performed after down-sampling read counts for protein-coding genes, to match those of lncRNAs (Methods). **D.** Distribution of the developmental stage in which maximum expression is observed, for protein-coding genes and lncRNAs that are significantly DE (FDR<0.01) in each organ, for the mouse. The percentages are computed with respect to the total number of DE genes in each organ and gene class.

As noted previously, between 59 and 82% of protein-coding genes are significantly differentially expressed (DE) among developmental stages, at a false discovery rate (FDR) below 1%, in each organ and species (Figure 4C, Supplementary Dataset 4). The proportions of DE lncRNAs are much lower in somatic organs, between 18 and 40%, but are similar in the testes, around 75% (Figure 4C). However, we suspected that this could be due to the low expression levels of this class of genes, as total read counts are known to affect the sensitivity of DE analyses (Anders and Huber 2010). Indeed, as previously observed, lncRNAs are expressed at much lower levels than protein-coding genes (Supplementary Figure 7). To control for this effect, we down-sampled the read counts observed for protein-coding genes, bringing them to the same average counts as lncRNAs but preserving relative gene abundance (Methods). Strikingly, when performing the DE analysis on this dataset, we observe higher proportions of DE loci for lncRNAs compared to protein-coding genes (Figure 4C). Moreover, the amplitude of expression variation among developmental stages are more important for lncRNAs than for protein-coding genes (Supplementary Figure 8A). This is expected given the lower lncRNA expression levels, which preclude detecting subtle expression shifts among time points. Finally, we observe that the developmental stage with maximum expression is generally different between protein-coding genes and lncRNAs, even when considering genes that are significantly DE among stages. For all organs, DE lncRNAs tend to show highest expression levels in the young and aged adults, while DE protein-coding genes are more homogeneously distributed among developmental stages (Figure 4D, Supplementary Figure 8B).

Similar conclusions are reached when performing DE analyses between consecutive time points (Supplementary Dataset 4, Supplementary Figure 6). For both protein-coding genes and lncRNAs, the strongest expression changes are observed between newborn and young adult individuals. Almost 10,000 lncRNAs are significantly up-regulated between newborn and young adult testes, confirming the strong enrichment for lncRNAs during spermatogenesis (Supplementary Dataset 4). We note that, as expected, the lowest numbers of DE genes are observed at the transition between young and aged adult organs. At this time-point, we observe more changes for the rat than for the mouse, potentially due to a higher proportion of immune cell infiltrates in rat aged organs. Genes associated with antigen processing and presentation tend to be expressed at higher levels in aged adults than in young adults, for mouse kidney, rat brain and liver (Supplementary Dataset 4).

### Stronger selective constraint on lncRNAs expressed earlier in development

We next analyzed the patterns of long-term evolutionary sequence conservation for lncRNAs, in conjunction with their spatio-temporal expression pattern (Supplementary Table 6). We used the PhastCons score (Siepel et al. 2005) across placental mammals (Casper et al. 2018), to assess the level of sequence conservation for various aspects of mouse lncRNAs: exonic promoter regions (defined as 1 kb regions upstream of the transcription start site, masking any exonic sequence within this region), splice sites (first and last two bases of the introns, for multi-exonic loci). As approximately 20% of lncRNAs overlap with exonic regions from other genes on the opposite strand (Supplementary Dataset 1), we masked exonic sequences from other genes before computing sequence conservation scores. As previously observed (Ponjavic et al. 2007; Haerty and Ponting 2013), the amount of exonic and splice site sequence conservation is much lower for lncRNAs than for protein-coding genes, but above the average conservation observed for intergenic regions (Figure 5A, C). In contrast, promoter sequence conservation levels are more comparable between protein-coding genes and lncRNAs (Figure 5B). The highest levels of conservation are observed for bidirectional promoters, shared with protein-coding genes (Figure 5B).

**Figure 5.**
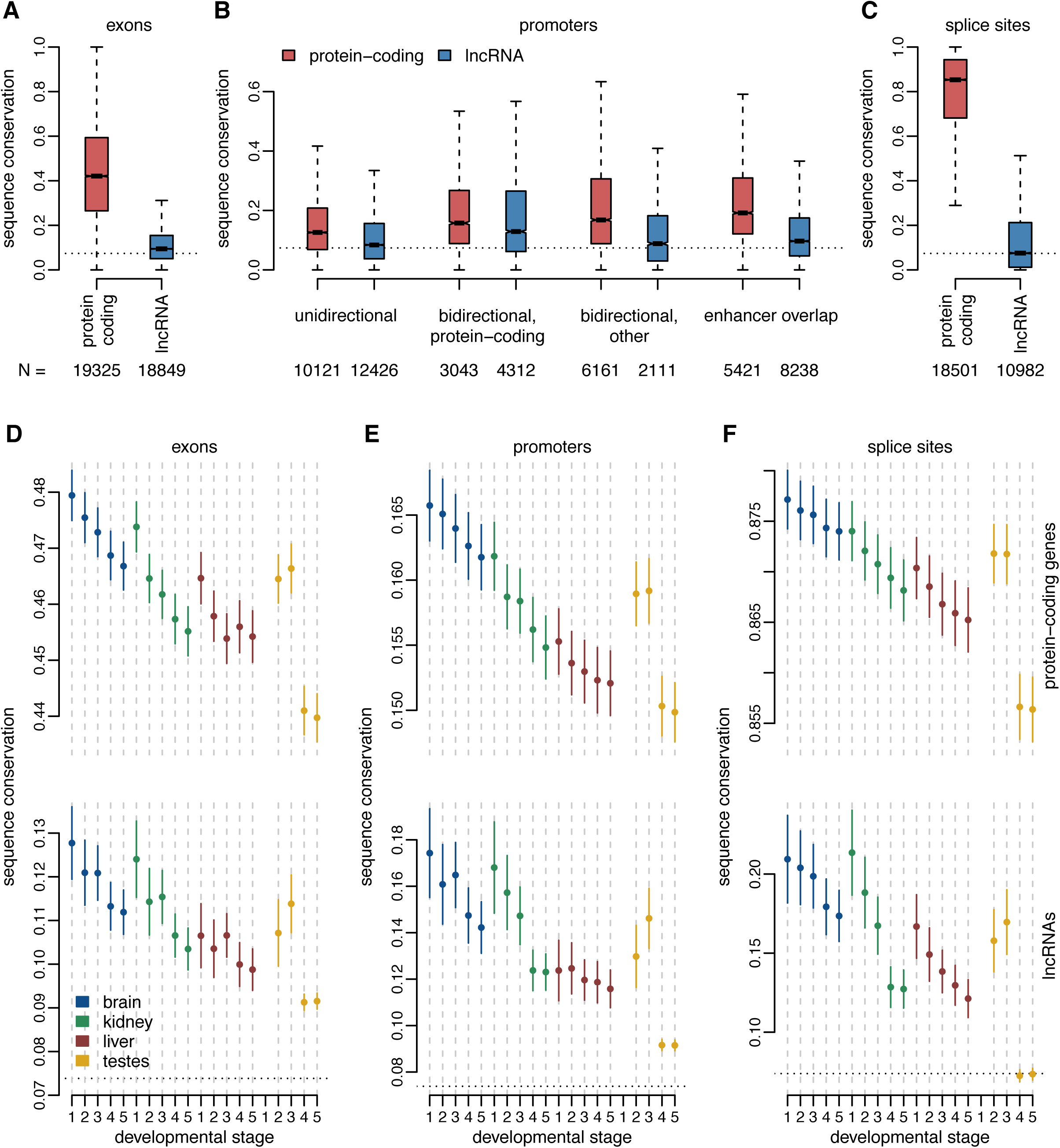
Increased levels of long-term sequence conservation for lncRNAs expressed early in development. **A.** Distribution of the sequence conservation score (PhasCons), for protein-coding and lncRNAs exonic regions, in the mouse. We used precomputed PhastCons score for placental mammals, downloaded from the UCSC Genome Browser. Exonic regions that overlap with exons from other genes were masked. Boxplot notches represent 95% confidence intervals of the medians. Numbers of analyzed genes are shown below the plot. The horizontal dotted line represents the average conservation score computed on intergenic regions, at least 5kb away from transcribed loci. **B.** Distribution of the sequence conservation score, for promoter regions (1kb upstream of transcription start sites) of mouse protein-coding genes and lncRNAs. Exonic sequences were masked in promoter regions. Genes are divided in different classes depending on their promoter type: unidirectional, bidirectional shared with protein-coding genes, bidirectional shared with non-coding genes, overlap with Encode-annotated enhancers. **C.** Distribution of the sequence conservation score, for splice sites (first and last two bases of each intron). **D.** Distribution of the sequence conservation score for protein-coding and lncRNAs exonic regions, for subsets of genes expressed above noise levels (TPM>=1) in each organ and developmental stage. Dots represent medians, vertical bars represent 95% confidence intervals. Numbers of analyzed genes are provided in Supplementary Table 5. **E.** Same as A, for promoter regions (1kb upstream of transcription start sites). Exonic sequences were masked before assessing conservation. **F.** Same as B, for splice sites (first and last two bases of each intron).

We next analyzed sets of protein-coding genes and lncRNAs that are expressed above noise levels (TPM>=1, averaged across all replicates) in each organ / developmental stage combination. For exonic sequences and splice site regions, the extent of sequence conservation is much lower for lncRNAs than for protein-coding genes, irrespective of the organ and developmental stage in which they are expressed (Figure 5D, F). For all examined regions and for both categories of genes, the spatio-temporal expression pattern is well correlated with the level of sequence conservation. Globally, sequence conservation is higher for genes that are expressed earlier in development than for genes expressed later in development, and is significantly higher for somatic organs than for adult and aged testes (Figure 5D-F). Interestingly, for genes that are highly expressed in mid-stage embryonic brain and kidney samples, the levels of promoter sequence conservation are higher for lncRNAs than for protein-coding genes (Figure 5E). However, this pattern is mainly due to those lncRNA loci that have bidirectional promoters, shared with protein-coding genes or with other non-coding loci (Supplementary Figure 9A-C).

Finally, we asked whether the highest level of evolutionary sequence conservation is seen at exons, promoter or splice site regions, for each lncRNA locus taken individually. We show that this pattern also depends on the organs and the developmental stages where the lncRNAs are expressed: for loci detected in somatic organs and in the developing testes, there is significantly higher conservation for the promoter and the splice sites than for exonic regions (Supplementary Figure 9D, E). However, for lncRNAs that are highly transcribed in the adult and aged testes (which constitutes the great majority of genes), this pattern is absent (Supplementary Figure 9D, E).

### Detection of homologous lncRNAs across species

Having investigated the patterns of long-term sequence conservation of mouse lncRNAs, we next sought to assess the conservation of lncRNA repertoires in mouse, rat and chicken. We detected lncRNA separately in each species, using only RNA-seq data and existing genome annotations, as previously suggested (Hezroni et al. 2015). We then searched for putative 1-to-1 orthologous lncRNAs between species using pre-computed whole-genome alignments as a guide (Methods), to increase the sensitivity of orthologous gene detection in the presence of rapid sequence evolution (Washietl et al. 2014). The orthologous lncRNA detection procedure involves several steps, including the identification of putative homologous (projected) loci across species, filtering to remove large-scale structural changes in the loci and intersection with predicted loci in the target species (Methods). As illustrated in Figure 6, for comparisons between rodents the extent of sequence divergence is low enough that more than 90% of 18,858 lncRNA loci are successfully projected from mouse to rat (Figure 6A, Supplementary Dataset 5). However, only 54% of projected loci have even weak levels of detectable transcription in the target species (at least 10 uniquely mapped reads). Only 26% of mouse lncRNA loci have predicted 1-to-1 orthologues in the rat, and only 15% are orthologous to confirmed lncRNA loci in the rat (Figure 6A, Supplementary Dataset 5). The 1,493 mouse lncRNAs that have non-lncRNAs orthologues in the rat are generally matched with loci discarded because of low read coverage, minimum exonic length or distance to protein-coding genes (Supplementary Dataset 5). Cases of lncRNA-protein-coding orthologues are rare at this evolutionary distance (Supplementary Dataset 5), and they may stem from gene classification errors. We note that, even when orthologous loci can be detected, lncRNA gene structures are highly divergent across species, in terms of exonic length or number of exons (Supplementary Figure 10).

**Figure 6.**
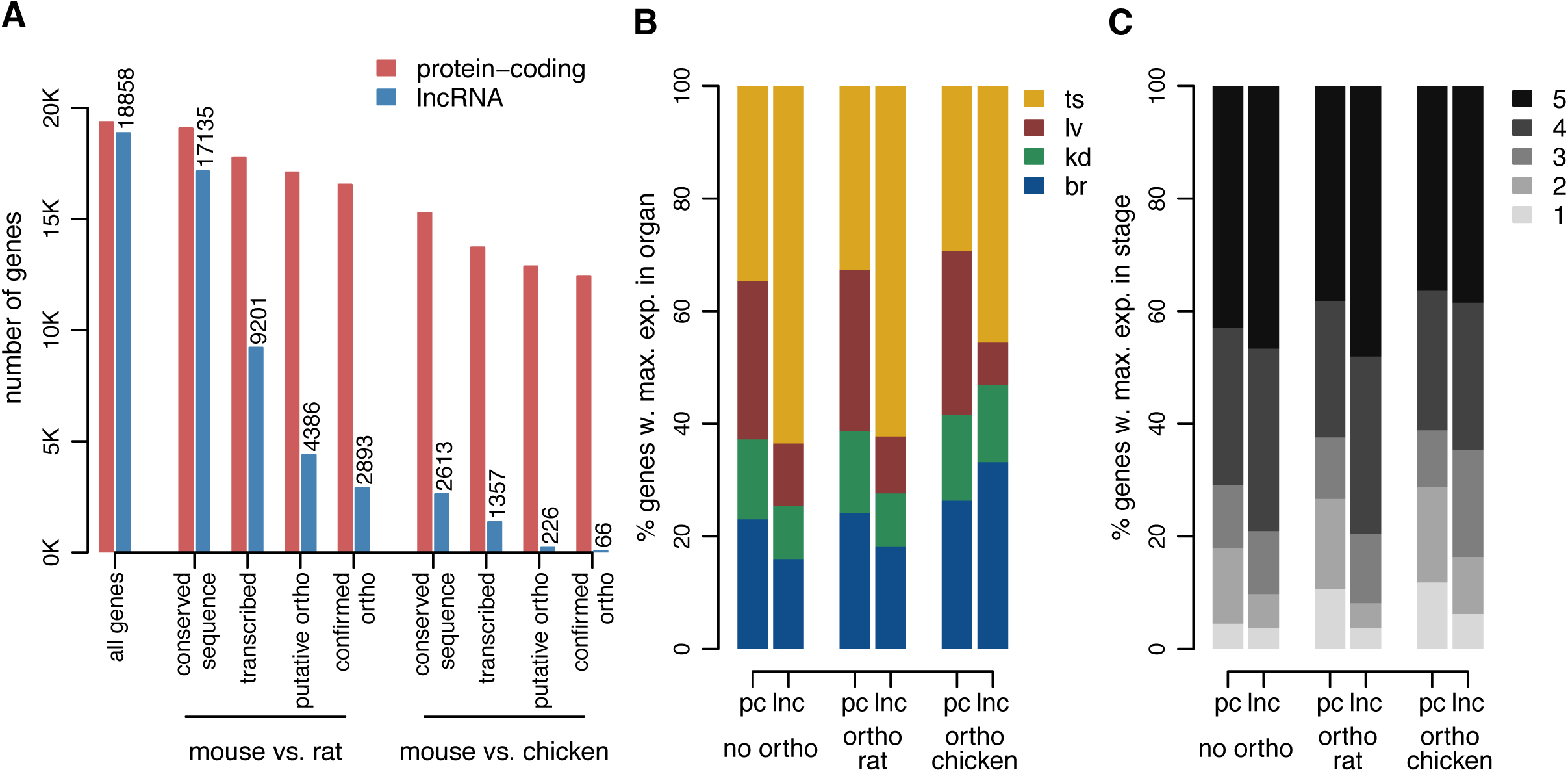
Orthologous lncRNA families for mouse, rat and chicken. **A.** Number of mouse protein-coding genes and lncRNAs in different classes of evolutionary conservation. From left to right: all loci (with TPM>=1 in at least one mouse sample), loci with conserved sequence in the rat, loci for which transcription could be detected (at least 10 unique reads) in predicted orthologous locus in the rat, loci with predicted 1-to-1 orthologues, loci for which the predicted orthologue belonged to the same class (protein-coding or lncRNA) in the rat, loci with conserved sequence in the chicken, loci for which transcription could be detected (at least 10 unique reads) in predicted orthologous locus in the chicken, loci with predicted 1-to-1 orthologues, loci for which the predicted orthologue belonged to the same class (protein-coding or lncRNA) in the chicken. We analyze 19,356 protein-coding genes and 18,858 candidate lncRNAs in the mouse. **B.** Distribution of the organ in which maximum expression is observed, for mouse protein-coding and lncRNA genes that have no orthologues in the rat or chicken, for genes with orthologues in the rat and for genes with orthologues in chicken. **C.** Same as B, for the distribution of the developmental stage in which maximum expression is observed.

At larger evolutionary distances, the rate of sequence evolution is the main factor hampering detection of orthologous lncRNAs. Only 2,613 (14%) of mouse lncRNAs could be projected on the chicken genome, and after subsequent filters we detect only 66 mouse – chicken lncRNA orthologues (Supplementary Dataset 5). We note that our lncRNA detection power is weaker for the chicken than for the rodents because of organ and developmental stage sampling, although we did strive to include RNA-seq data from adult organs in the lncRNA detection process (Methods, Supplementary Table 2). Conserved lncRNAs differ from non-conserved lncRNAs in terms of expression patterns. While only subtle differences can be observed when comparing mouse-rat orthologous lncRNAs to the mouse-specific lncRNA set, lncRNAs that are conserved across rodents and chicken (Supplementary Table 7) are strongly enriched in somatic organs and early developmental stages (Figure 6B,C).

### Global patterns of lncRNA expression across species, organs and developmental stages

We next assessed the global patterns of expression variation across species, organs and developmental stages, for predicted mouse – rat lncRNA orthologues (Supplementary Dataset 6). As for protein-coding genes, the main source of variability in a PCA performed on lncRNA expression levels is the difference between adult and aged testes and the other samples (Figure 7A). However, for lncRNAs, samples cluster according to the species of origin already on the second factorial axis (11% explained variance), thus confirming that lncRNA expression patterns evolve rapidly. Overall, differences between organs and developmental stages are less striking for lncRNAs, compared to the variation stemming from the species factor (Figure 7A). This pattern is also visible on a hierarchical clustering analysis (performed on distances derived from Spearman’s correlation coefficient): in contrast with what is observed for protein-coding genes, for lncRNAs samples generally cluster by species, with the exception of adult and aged testes which are robustly grouped (Figure 7B).

**Figure 7.**
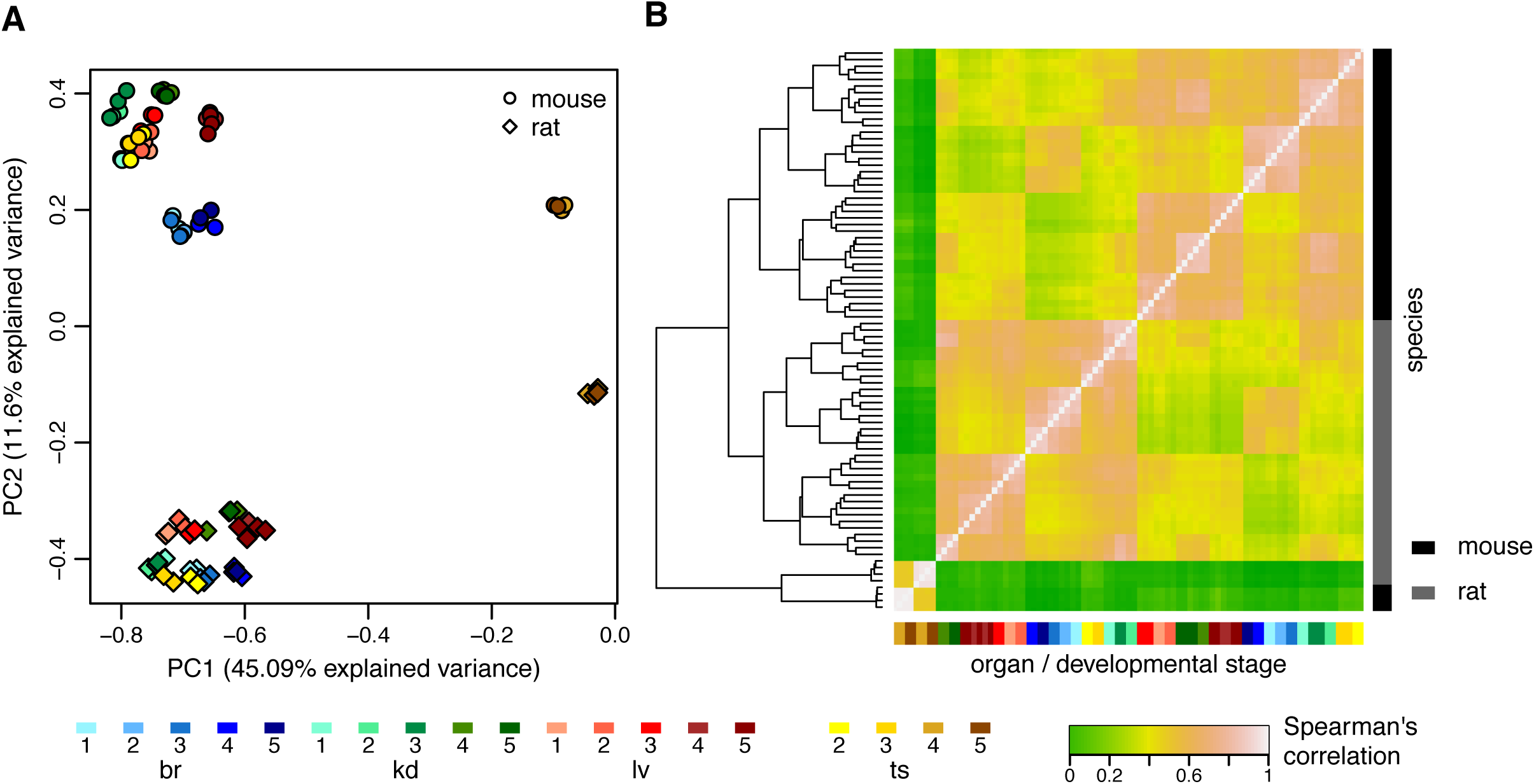
Global comparison of lncRNA expression patterns across species. **A.** First factorial map of a principal component analysis, performed on log2-transformed TPM values, for 2,893 orthologous lncRNAs between mouse and rat. Colors represent different organs and developmental stages, point types represent species. **B.** Hierarchical clustering, performed on a distance matrix derived from Spearman correlations between pairs of samples, for 2,893 orthologous lncRNAs between mouse and rat. Organ and developmental stages are shown below the heatmap. Species of origin is shown on the right. Sample clustering is shown on the left.

The higher rates of lncRNA expression evolution are also visible when analyzing within-species variations, through comparisons across biological replicates (Figure 8A). Both within-species and between-species comparisons are affected by technical biases, such as the low expression levels of lncRNAs, which hampers accurate expression estimates. To partially account for this effect, we sought to measure the global extent of gene expression conservation by contrasting between-species and within-species variations. Briefly, we constructed an expression conservation index by dividing the between-species and the within-species Spearman’s correlation coefficient, computed on all genes from a category, for a given organ/developmental stage combination (Methods). The resulting expression conservation values are very high for protein-coding genes, in particular for the brain and the mid-stage embryonic kidney. However, there is significant less conservation between species for the adult and aged testes (Figure 8B). For lncRNAs, expression conservation values are much lower than those observed for protein-coding genes, with strikingly low values for adult and aged testes (Figure 8C).

**Figure 8.**
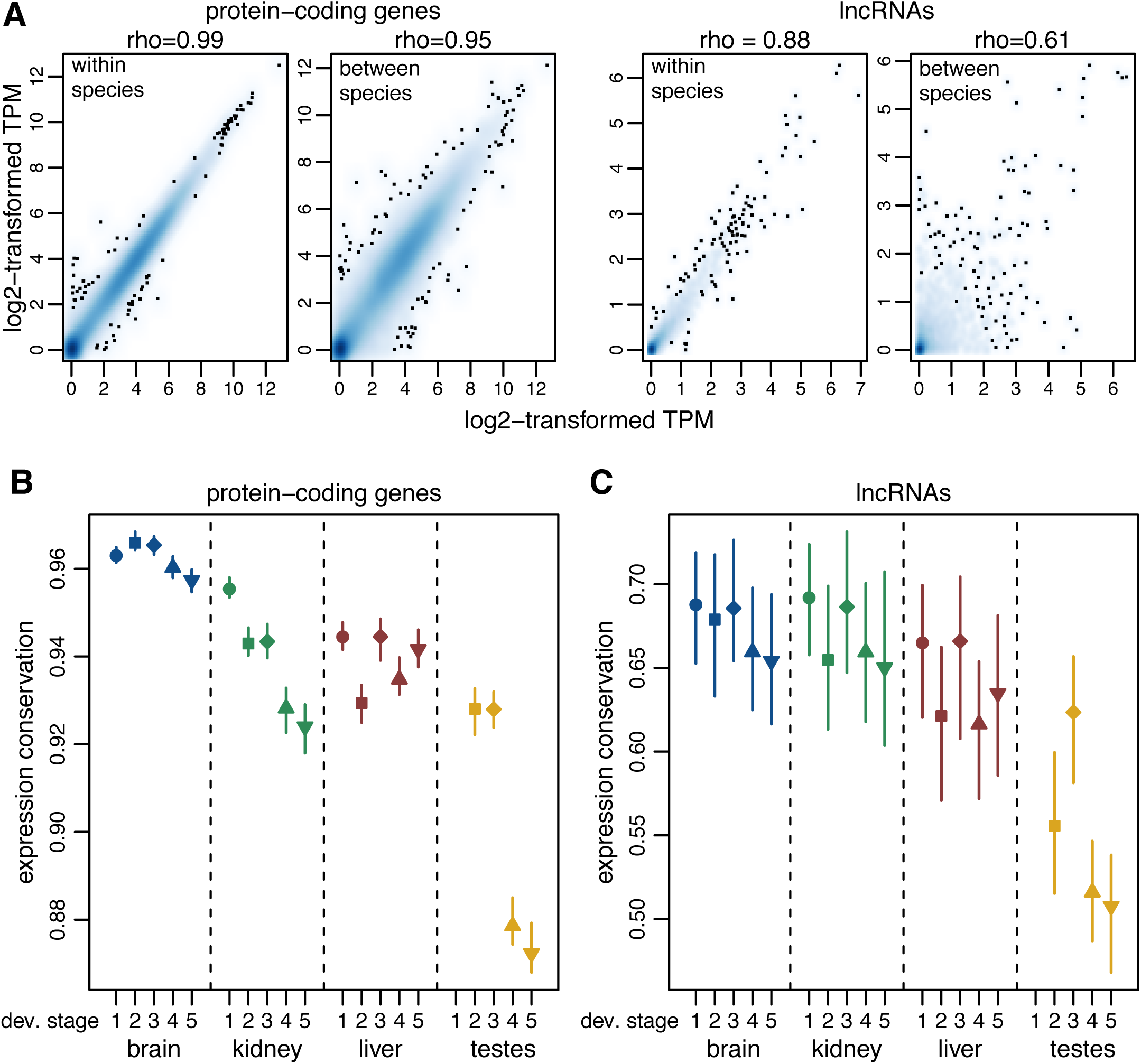
Global estimates of expression conservation across organs and developmental stages. **A.** Example of between-species and within-species variation of expression levels, for protein-coding genes (left) and lncRNAs (right), for orthologous genes between mouse and rat, for the mid-stage embryonic brain. Spearman’s correlation coefficients (rho) are shown above each plot. We show a smoothed color density representation of the scatterplots, obtained through a (2D) kernel density estimate (smoothScatter function in R). **B.** Expression conservation index, defined as the ratio of the between-species and the within-species expression level correlation coefficients, for protein-coding genes, for each organ and developmental stage. The vertical segments represent minimum and maximum values obtained from 100 bootstrap replicates. We analyzed 15,931 pairs of orthologous genes between mouse and rat. **C.** Same as B, for lncRNAs. We analyzed 2,893 orthologous mouse and rat lncRNAs.

### Evolutionary divergence of individual lncRNA expression profiles

For the mouse and rat, we could delve deeper into the evolutionary comparison of lncRNA expression patterns, by asking whether variations among developmental stages are shared between species, for individual lncRNAs. We used models from the DESeq2 (Love et al. 2014) package to detect differential gene expression among developmental stages, independently for each species and organ (Supplementary Dataset 4, Methods). We focused on orthologous lncRNAs that are significantly differentially expressed (FDR<0.01) in both mouse and rat, and compared their patterns of expression variations among developmental stages. Several hundreds of orthologous lncRNAs are DE in both mouse and rat, in each organ (Figure 9). These lncRNAs show parallel patterns of variation among developmental stages in mouse and rat, for all organs (Figure 9). Indeed, the developmental stage with maximum expression is generally coherent across species (Figure 9A). As previously observed, few genes are differentially expressed between young and aged adults (Supplementary Figure 6), hence these two developmental stages are mixed in the comparative DE analysis. We clustered the relative expression profiles of shared DE lncRNAs across species using the k-means algorithm (Methods). These patterns further illustrate the similarity of the relative expression profiles among species (Figure 9B-E). Interestingly, although temporal expression profiles are likewise similar between mouse and rat for DE protein-coding genes (Supplementary Figure 11), almost 25% of shared DE protein-coding genes show different trends for mouse and rat in the testes (Supplementary Figure 11). These sets of genes do not show any strong functional enrichment (Supplementary Dataset 4). This pattern confirms previous reports of rapid expression evolution in the adult testes (Brawand et al. 2011), and extends them by showing that patterns of variations among developmental stages are often species-specific in the testes, for protein-coding genes.

**Figure 9.**
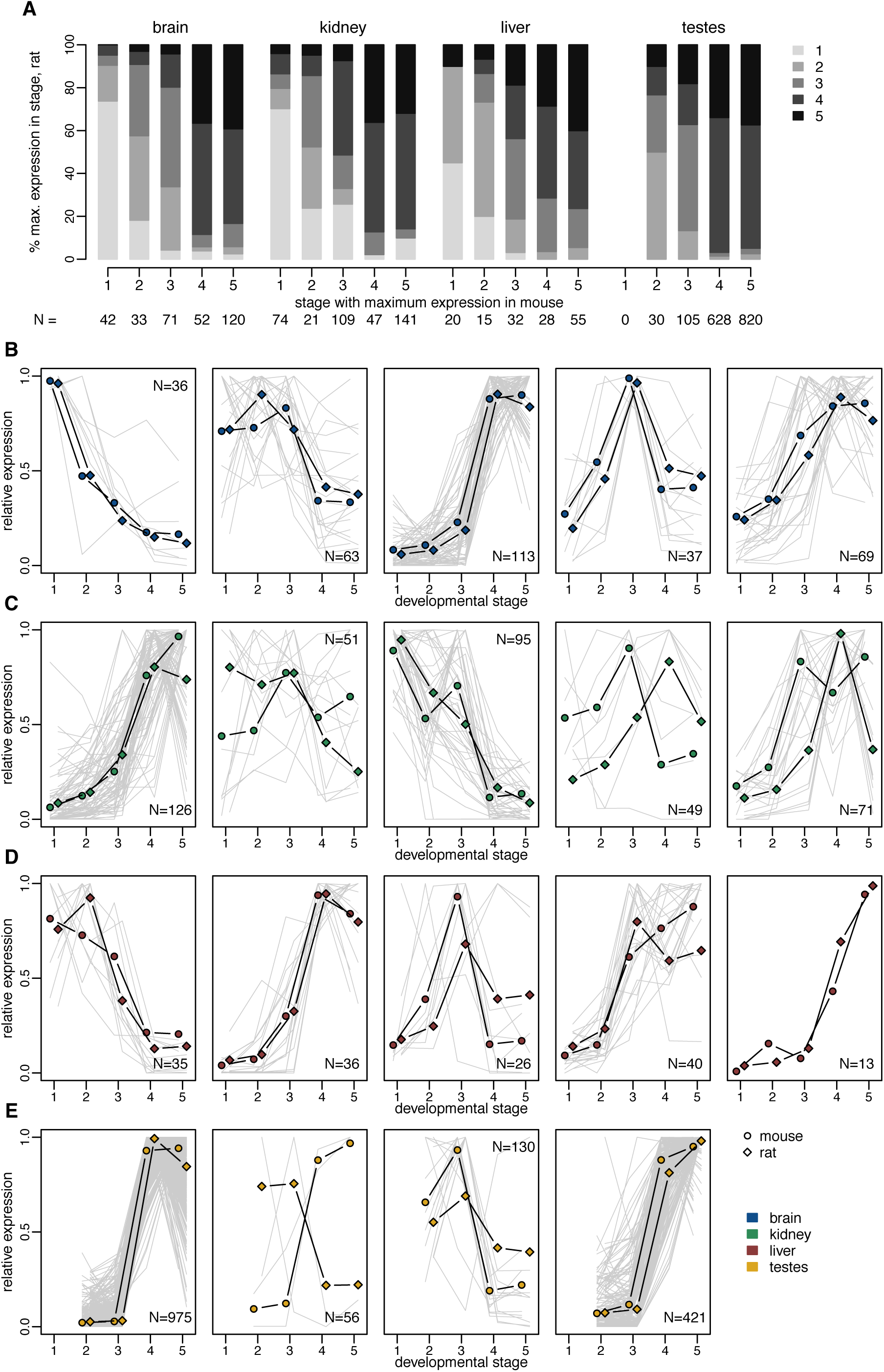
Conservation of developmental expression patterns of differentially expressed lncRNAs. **A.** Comparison of the developmental stage in which maximum expression is observed, for orthologous lncRNAs that are significantly differentially expressed (FDR<0.01) among developmental stages, for both mouse and rat. Genes are divided based on the developmental stage where maximum expression is observed in mouse organs (X-axis). The Y axis represents the percentage of orthologous genes that reach maximum expression in each developmental stage, in the rat. Numbers of analyzed genes are shown below the plot. **B.** Expression profiles of orthologous lncRNAs that are significantly differentially expressed (FDR<0.01) among developmental stages, for both mouse and rat, in the brain. TPM values were averaged across replicates and normalized by dividing by the maximum, for each species. The resulting relative expression profiles were combined across species and clustered with the K-means algorithm. The average profiles of the genes belonging to each cluster are shown. Gray lines represent profiles of individual genes from a cluster. Numbers of genes in each cluster are shown in the plot. **C.** Same as B, for the kidney. **D.** Same as B, for the liver. **E.** Same as B, for the testes. For this organ, we searched for only 4 clusters with the K-means algorithm.

To further quantify lncRNA expression profile differences among species, we measured the amount of expression divergence as the Euclidean distance between relative expression profiles (average TPM values across biological replicates, normalized by dividing by the sum of all values for a gene, for each species), for mouse and rat orthologues (Methods, Supplementary Dataset 7, Supplementary Table 8). The resulting expression divergence values correlate negatively with the average expression level (Figure 10A), as expected given that abundance estimation is less reliable for weakly expressed genes. While the raw expression divergence values are significantly higher for lncRNAs than for protein-coding genes (Figure 10B), this is largely due to the low lncRNA expression levels. Indeed, the effect disappears when analyzing the residual expression divergence after regressing the mean expression level (Figure 10C). These patterns remain true when analyzing separately protein-coding and lncRNAs with different types of promoters, bidirectional or unidirectional (Supplementary Figure 12A). For lncRNAs, we also observe a weak negative correlation between expression divergence and the extent of gene structure conservation (Figure 10D). We measured the relative contribution of each organ/developmental stage to the expression divergence estimate (Figure 10E). For both protein-coding genes and lncRNAs, by far the highest contributors are the young adult and aged testes samples, which are responsible for almost 30% of the lncRNA expression divergence (Figure 10E). This is visible in the expression patterns of the 2 protein-coding and lncRNA genes with the highest residual expression divergence: the lncRNA expression divergence is mostly due to changes in adult testes, while more complex expression pattern changes seem to have occurred for the protein-coding genes (Supplementary Figure 12). The most divergent protein-coding genes are enriched in functions related to immunity (Supplementary Dataset 7), suggesting that differences in immune cell infiltrates among species could be responsible for these extreme cases of expression pattern divergence.

**Figure 10.**
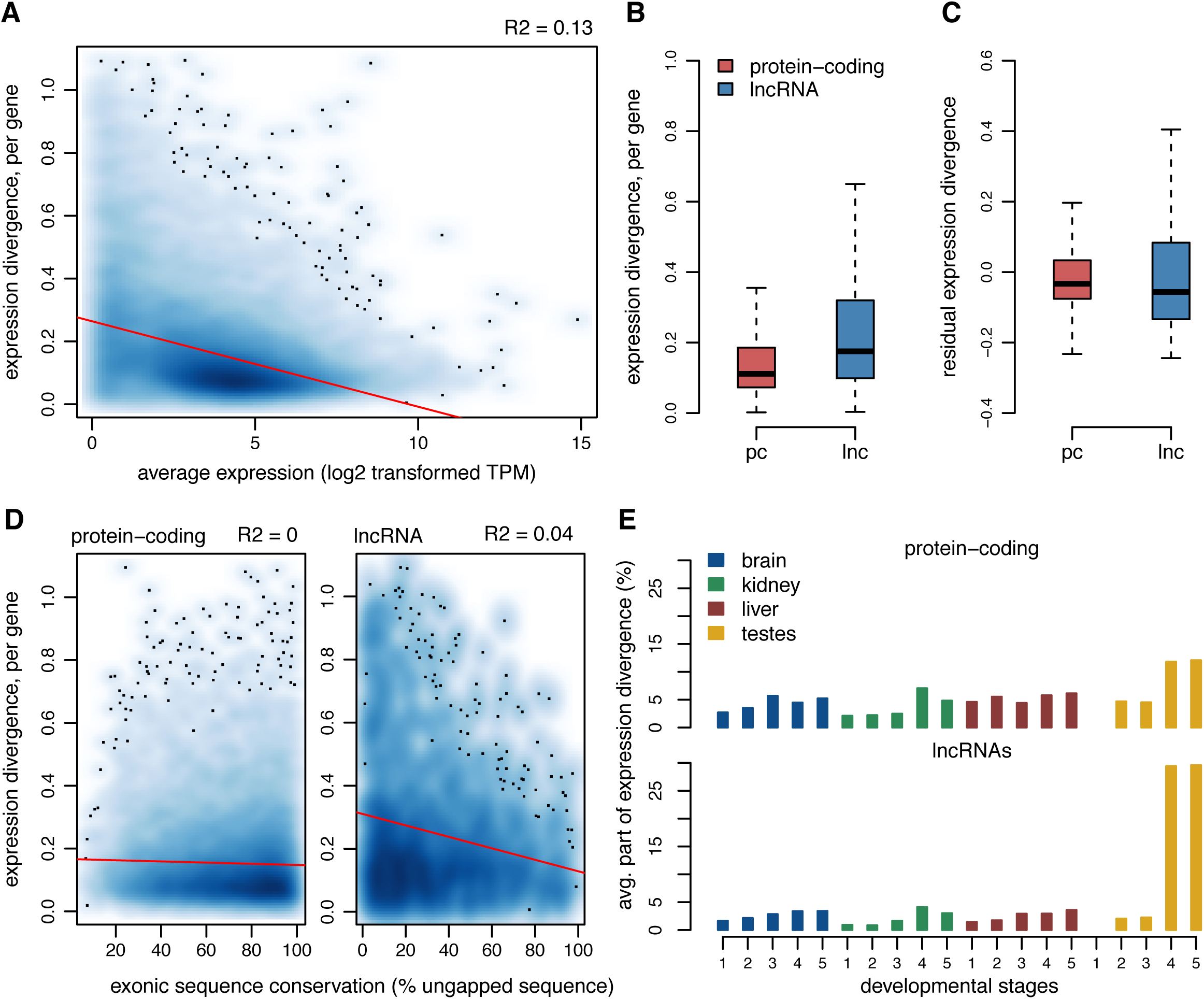
*Per*-gene estimates of expression pattern divergence between species. **A.** Relationship between the per-gene expression divergence measure (Euclidean distance of relative expression profiles among organs/stages, between mouse and rat), and the average expression values (log2-transformed TPM) across all mouse and rat samples. We show a smoothed color density representation of the scatterplots, obtained through a (2D) kernel density estimate (smoothScatter function in R). Red line: linear regression. **B.** Distribution of the expression divergence value for all protein-coding and lncRNA genes with predicted 1-to-1 orthologues in mouse and rat. **C.** Distribution of the residual expression divergence values, after regressing the average expression level, for protein-coding genes and lncRNAs. **D.** Relationship between expression divergence and exonic sequence conservation (% exonic sequence aligned without gaps between mouse and rat), for protein-coding genes and lncRNAs. **E.** Average contribution of each organ/developmental stage combination to expression divergence, for protein-coding genes and lncRNAs.

### Candidate species-specific lncRNAs

We next sought to investigate the most extreme cases of expression divergence: situations where expression can be robustly detected in one species, but not in the other one, despite the presence of almost perfect sequence alignment (Methods). We selected lncRNA loci that were supported by at least 100 uniquely mapped reads in one species, with no reads detected in the predicted homologous region in the other species. With this convention, we obtain 1,041 candidate mouse-specific and 1,646 candidate rat-specific loci (Supplementary Dataset 8). These lists include striking examples, such as the region downstream of the *Fzd4* protein-coding gene, which contains a mouse-specific and a rat-specific lncRNA candidate, each perfectly aligned in the other species (Supplementary Figure 13A). We could not identify any differential transcription factor binding or transposable element enrichment in the promoters of these species-specific lncRNAs (data not shown). Interestingly however, they are increasingly associated with predicted expression enhancers (Supplementary Figure 14). While the evolutionary and mechanistic origin of these lncRNAs is still mysterious, we could confirm that their presence is associated with increased expression divergence in the neighboring genes. To test this, we selected species-specific and orthologous lncRNAs that are transcribed from bidirectional promoters shared with protein-coding genes, and evaluated the expression divergence of their protein-coding neighbors (Supplementary Figure 13B,C). Though the difference is subtle, genes that are close to species-specific lncRNAs have significantly higher expression divergence than the ones that have conserved lncRNA neighbors, even after correcting for expression levels (Wilcoxon test, p-value < 10-3). It thus seems that expression changes that led to the species-specific lncRNA transcription extend beyond the lncRNA locus and affect the neighboring genes, as previously proposed (Kutter et al. 2012).

## Discussion

### Assessing lncRNA functionality: current challenges and insights from evolutionary approaches

More than a decade after the publication of the first genome-wide lncRNA datasets (Guttman et al. 2009; Khalil et al. 2009), the debate regarding their functionality is still not settled. While experimental assessments of lncRNA functions are rapidly accumulating, they are lagging behind the exponential increase of RNA sequencing datasets, each one revealing thousands of previously unreported noncoding transcripts (Pertea et al. 2018). There is thus a need to define biologically relevant criteria to prioritize lncRNAs for experimental investigation. Furthermore*, in vivo* tests of lncRNA functions need to be carefully designed to account for ubiquitous confounding factors, such as the presence of overlapping regulatory elements at lncRNA loci (Bassett et al. 2014). Another challenge is the fact that some lncRNA loci undoubtedly have “unconventional” biological functions, that require for example the presence of a transcription and splicing at a given genomic location, independently of the lncRNA molecule that is produced (Latos et al. 2012; Engreitz et al. 2016).

Evolutionary approaches can provide important tools to assess biological functionality (Haerty and Ponting 2014), and they have been already successfully applied to lncRNAs. Although only a few large-scale comparative transcriptomics studies are available so far for vertebrate lncRNAs (Kutter et al. 2012; Washietl et al. 2014; Hezroni et al. 2015; Necsulea et al. 2014), they all agree that lncRNAs evolve rapidly in terms of primary sequence, exon-intron structure and expression patterns, indicating that there is little selective constraint and thus little functionality for these loci. However, these studies have all focused on lncRNAs detected in adult organs. We hypothesized that lncRNAs expressed during embryogenesis are enriched in functional loci, as suggested by the increasing number of lncRNAs with proposed roles in development (Rinn et al. 2007; Sauvageau et al. 2013; Grote et al. 2013; Grote and Herrmann 2015). To test this hypothesis, we performed a multi-dimensional comparative transcriptomics analysis, following lncRNA and protein-coding gene expression patterns across species, organs and developmental stages.

### Comparability of transcriptomes across species, organs and developmental stages

One of main concerns was to ensure that our transcriptome collection provides a good representation of the changes in cellular composition and physiological functions that occur during major organ development, and that the resulting patterns are comparable across diverse vertebrate species. Our analyses of cell-type specific gene markers, derived from single-cell transcriptomics analyses (Tabula Muris Consortium 2018; Green et al. 2018), confirms that this is indeed the case, as we see expected expression patterns in our transcriptome collection. Likewise, we observed due enrichment of biological processes in the sets of genes that are specifically expressed in each organ/developmental stage combination. Furthermore, we showed that protein-coding gene expression profiles across major organs and developmental stages are well conserved among species, even at large evolutionary distances. Although differences among rodents and chicken are visible when analyzing the full set of orthologous protein-coding genes, we find that the expression profiles of genes that are known to be implicated in embryonic development and in gene expression regulation processes are highly conserved among species (Figure 2). Our transcriptome collection thus enables detection of ancestral developmental expression patterns, shared across amniote species, for key players in developmental regulatory networks.

### Spatio-temporal lncRNA expression patterns

Our first major observation is that lncRNAs are overwhelmingly expressed in the adult and aged testes, in agreement with previous data (Soumillon et al. 2013). Their relative depletion in embryonic and newborn testes reinforces the association between lncRNA production and spermatogenesis, in accord with the hypothesis that the particular chromatin environment during spermatogenesis is a driver for promiscuous, non-functional transcription (Kaessmann 2010; Soumillon et al. 2013). Interestingly, we show that lncRNAs are significantly differentially expressed among developmental stages, at least as frequently as protein-coding genes, after correcting for their lower expression levels. However, in contrast with protein-coding genes, the majority of lncRNAs reach their highest expression levels in adult rather than in developing organs. As requirements for tight gene expression control are undoubtedly higher during embryonic development (Ben-Tabou de-Leon and Davidson 2007), an explanation for the relative lncRNA depletion in embryonic and newborn transcriptomes is that transcriptional noise is more efficiently blocked during the early stages of development. Differences in cellular composition heterogeneity may also be part of the explanation. Expression analyses of cell-type specific markers suggest that adult and aged organ transcriptomes may be a mix of more diverse cell types, notably including substantial immune cell infiltrates. A higher cell type diversity may explain the increased abundance of lncRNAs in adult and aged organs, especially given that lncRNAs are thought to be cell-type specific (Liu et al. 2016).

### Functionally constrained lncRNAs are enriched in developmental transcriptomes

We show that, for those lncRNAs that are expressed above noise levels (TPM>=1) in somatic organs and in the earlier developmental stages, there is a higher proportion of functionally constrained loci than in testes-expressed lncRNAs. Strikingly, we find that the level of long-term sequence conservation for lncRNA promoter regions is higher than the one observed for protein-coding promoters, when we analyze genes that are robustly expressed (TPM>=1) in embryonic brain and kidney. Furthermore, we show that lncRNAs that are expressed in somatic organs and in the developing testes differ from those expressed in the adult testes not only in terms of overall levels of sequence conservation, but also with respect to the regions of the lncRNA loci that are under selective constraint. Thus, for lncRNAs that are expressed in somatic organs and in the developing testes, there is significantly more evolutionary constraint on promoter and splice site sequences than on exonic regions, while these patterns are not seen for the bulk of lncRNAs, expressed in adult and aged testes. We are thus able to modulate previous reports of increased constraint on splicing regulatory regions in mammalian lncRNAs (Schüler et al. 2014; Haerty and Ponting 2015), by showing that this pattern is specifically seen in lncRNAs that are expressed in somatic organs and in the developing testes.

These results are also in agreement with a series of recent findings, suggesting that at many lncRNA loci, biological function may reside in the presence of additional non-coding regulatory elements at the lncRNA promoter rather than in the production of a specific transcript (Engreitz et al. 2016; Groff et al. 2016). While the elevated sequence conservation at splicing regulatory signals could in principle indicate that the production of a specific mature lncRNA molecule is required, we note that splicing of lncRNA transcripts was recently proposed to affect the expression of neighboring protein-coding genes (Engreitz et al. 2016). Thus, while there is evidence for increased functionality for those lncRNA loci that are detected in developmental transcriptomes or in adult somatic organs, our sequence conservation analyses suggest that their biological functions may be carried out in an RNA-independent manner, as exonic sequences are under less constraint than promoter or splice site regions.

### Evolutionary divergence of spatio-temporal expression profiles for lncRNAs

We previously established that lncRNA expression patterns evolve rapidly across species in adult organs. Here, we show that this rapid evolution of lncRNA expression is not restricted to adult and aged individuals, but is also true for embryonic and newborn developmental stages. Expression patterns comparisons across species, organs and developmental stages are dominated by differences between species for lncRNAs, while similarities between organs and developmental stages are predominant for protein-coding genes, even across distantly related species. We assessed the extent of expression level conservation by contrasting between-species and within-species expression variations and we showed that lncRNAs have significantly lower levels of conservation than protein-coding genes, for all organs and developmental stages. However, lncRNA expression is significantly more conserved in somatic organs and in early embryonic stages than in the adult testes. Moreover, when orthologous lncRNAs are differentially expressed among developmental stages in both mouse and rat, they generally show parallel profiles of expression variation in both species. This observation is compatible with previous reports indicating that lncRNA expression is often cell type-specific (Liu et al. 2016). The differentially expressed lncRNAs, shared across mouse and rat, may be specific of cell types that change their relative abundance in whole-organ transcriptomes with developmental time.

Interestingly, when we evaluate expression divergence individually for each orthologous gene pair, and when we correct for the lower lncRNA expression levels, we find that lncRNAs are comparable with protein-coding genes, on average. This observation indicates that much of the between-species differences in lncRNA expression patterns may be due to the low expression levels of lncRNAs. It is not clear however whether this purely an indication of technical biases, that hamper accurate expression estimation for lowly expressed lncRNAs, or whether the low lncRNA expression levels can be interpreted as a sign that these transcripts are non-functional. For those lncRNAs that are cell type-specific, low expression levels in whole organ transcriptomes are expected. This question may be directly addressed in the near future, as single-cell transcriptome assays become more sensitive and allow investigation of lncRNA expression patterns (Liu et al. 2016).

### Candidate species-specific lncRNAs

Finally, we analyzed extreme cases of expression divergence between species, namely situations where transcription can be robustly detected in one species but not in the other, despite the presence of good sequence conservation. We identify more than a thousand candidate species-specific lncRNAs, in both mouse and rat. Interestingly, we observe that candidate mouse-specific lncRNAs are more frequently transcribed from enhancers than lncRNAs conserved between mouse and rat. This observation is consistent with previous reports that enhancers and enhancer-associated lncRNAs evolve rapidly (Villar et al. 2015; Marques et al. 2013). The genetic basis of these extreme transcription pattern changes is still not elucidated, and deserves further detailed investigations. Nevertheless, we show that these lncRNA expression patterns do not occur in an isolated manner. When such species-specific transcription was detected at protein-coding genes bidirectional promoters, the neighboring protein-coding genes also showed increased expression divergence, compared to genes that are transcribed from conserved lncRNA promoters. This observation is compatible with previous reports that lncRNA turnover is associated with changes in neighboring gene expression (Kutter et al. 2012). While lncRNAs changes may be directly affecting gene expression, it is also possible that a common mechanism affects both lncRNAs and protein-coding genes transcribed from bidirectional promoters.

## Conclusions

Our comparative transcriptomics approach confirms the established finding that lncRNAs repertoires, sequences and expression patterns evolve rapidly across species, and shows that the accelerated rates of lncRNA evolution are also seen in developmental transcriptomes. These observations are consistent with the hypothesis that the majority of lncRNAs (or at least of those detected with sensitive transcriptome sequencing approaches, in particular in the adult testes) may be non-functional. However, we are able to modulate this conclusion, by showing that there are increased levels of functional constraint on lncRNAs expressed during embryonic development, in particular in the developing brain and kidney. These increased levels of constraint apply to all analyzed aspects of lncRNAs, including sequence conservation for exons, promoter and splice sites, but also expression pattern conservation. For many of these loci, biological function may be RNA-independent, as the highest levels of selective constraint are observed on promoter regions and on splice signals, rather than on lncRNA exonic sequences. Our results are thus compatible with unconventional, RNA-independent functions for lncRNAs expressed during embryonic development.

## Methods

### Biological sample collection

We collected samples from three species (mouse C57BL/6J strain, rat Wistar strain and chicken White Leghorn strain), four organs (brain, kidney, liver and testes) and five developmental stages (including two embryonic stages, newborn, young and aged adult individuals). We sampled the following stages in the mouse: embryonic day post-conception (dpc) 13.5 (E13.5 dpc, hereafter mid-stage embryo); E17 to E17.5 dpc (late embryo); post-natal day 1 to 2 (newborn); young adult (8-10 weeks old); aged adult (24 months old). For the rat, we sampled the following stages: E15 dpc (mid-stage embryo); E18.5 to E19 dpc (late embryo); post-natal day 1 to 2 (newborn); young adult (8-10 weeks old); aged adult (24 months, with the exception of kidney samples and two of four liver samples, derived from 12 months old individuals). The embryonic and neonatal developmental stages were selected for maximum comparability based on Carnegie stage criteria (Theiler 1989). For chicken, we collected samples from Hamburger-Hamilton stages 31 and 36, hereafter termed mid-stage and late embryo. We selected these two stages for comparability with the two embryonic stages in mouse and rat (Hamburger and Hamilton 1951). In general, each sample corresponds to one individual, except for mouse and rat mid-stage embryonic kidney, for which tissue from several embryos was pooled prior to RNA extraction. For adult and aged organs, multiple tissue pieces from the same individual were pooled and homogenized prior to RNA extraction. For brain dissection, we sampled the cerebral cortex. For mouse and rat samples, with the exception of the mid-stage embryonic kidney, individuals were genotyped and males were selected for RNA extraction. Between two and four biological replicates were obtained for each species/organ/stage combination, amounting to 97 samples in total (Supplementary Table 1).

### RNA-seq library preparation and sequencing

We performed RNA extractions using RNeasy Plus Mini kit from Qiagen. RNA quality was assessed using the Agilent 2100 Bioanalyzer. Sequencing libraries were produced using the Illumina TruSeq stranded mRNA protocol with polyA selection, and sequenced as 101 base pairs (bp) single-end reads, at the Genomics Platform of iGE3 and the University of Geneva (https://ige3.genomics.unige.ch/).

### Additional RNA-seq data

To improve detection power for lowly expressed lncRNAs, we complemented our RNA-seq collection with samples generated with the same technology for Brown Norway rat adult organs (Cortez et al. 2014). We added data generated by the Chickspress project (http://geneatlas.arl.arizona.edu/) for adult chicken (red jungle fowl strain UCD001) organs, as well as for embryonic chicken (White Leghorn) organs from two publications (Uebbing et al. 2015; Ayers et al. 2013). Almost all samples were strand-specific, except the chicken adult organs and early embryonic testes. As the data were not perfectly comparable with our own in terms of library preparation and animal strains, the additional rat and chicken samples were only used to increase lncRNA detection sensitivity.

### RNA-seq data processing

We used HISAT2 (Kim et al. 2015) release 2.0.5 to align the RNA-seq data on reference genomes. The genome sequences (assembly versions mm10/GRCm38, rn6/Rnor_6.0 and galGal5/Gallus_gallus-5.0) were downloaded from the Ensembl database (Cunningham et al. 2019). Genome indexes were built using only genome sequence information. To improve detection sensitivity, at the alignment step we provided known splice junction coordinates extracted from Ensembl. We set the maximum intron length for splice junction detection at 1 million base pairs (Mb). To verify the strandedness of the RNA-seq data, we analyzed spliced reads that spanned introns with canonical (GT-AG or GC-AG) splice sites and compared the strand inferred based on the splice site with the one assigned based on the library preparation protocol (Supplementary Table 1). Finally, to estimate the mappability of each genomic region, we generated error-free artificial RNA-seq reads (single-end, 101 bp long, with 5 bp distance between consecutive read starts) from the genome sequence and realigned them to the genome with the same HISAT2 parameters. Regions for which the corresponding reads could be aligned unambiguously were considered “mappable”; the remaining regions were said to be “unmappable”.

### Transcript assembly and filtering

We assembled transcripts for each sample using StringTie (Pertea et al. 2015), release 1.3.5, based on read alignments obtained with HISAT2. We provided genome annotations from Ensembl release 94 as a guide for transcript assembly. We filtered Ensembl annotations to remove transcripts that spanned a genomic length above 2.5 Mb. For protein-coding genes, we kept only protein-coding transcripts, discarding isoforms annotated as “retained_intron”, “processed_transcript” etc. We set the minimum exonic length at 150 bp, the minimum anchor length for splice junctions at 8bp and the minimum isoform fraction at 0.05. We compared the resulting assembled transcripts with Ensembl annotations and we discarded read-through transcripts, defined as overlapping with multiple multi-exonic Ensembl-annotated genes. For strand-specific samples, we discarded transcripts for which the ratio of sense to antisense unique read coverage was below 0.01. We discarded multi-exonic transcripts that were not supported by splice junctions with correctly assigned strands. The filtered transcripts obtained for each sample were assembled into a single dataset *per* species using the merge option in StringTie. For increased sensitivity, we removed the minimum FPKM and TPM thresholds for transcript inclusion. We constructed a combined annotation dataset, starting with Ensembl annotations, to which we added newly-assembled transcripts that had no exonic overlap with Ensembl genes. We also included newly-annotated isoforms for known genes if they had exonic overlap with exactly one Ensembl gene, thus discarding potential read-through transcripts or gene fusions.

### Protein-coding potential of assembled transcripts

To determine whether the newly assembled transcripts were protein-coding or non-coding, we mainly relied on the codon substitution frequency (CSF) score (Lin et al. 2007). As in a previous publication (Necsulea et al. 2014) we scanned whole genome alignments and computed CSF scores in 75 bp sliding windows moving with a 3 bp step. We used pre-computed alignments downloaded from the UCSC Genome Browser (Casper et al. 2018), including the alignment between the mouse genome and 59 other vertebrates (for mouse classification), between the human genome and 99 other vertebrates (for rat and chicken classification) and between the rat genome and 19 other vertebrates (for rat classification). For each window, we computed the score in each of the 6 possible reading frames and extracted the maximum score for each strand. We considered that transcripts are protein-coding if they overlapped with positive CSF scores on at least 150 bp. As positive CSF scores may also appear on the antisense strand of protein-coding regions due to the partial strand-symmetry of the genetic code, in this analysis we considered only exonic regions that did not overlap with other genes. In addition, we searched for sequence similarity between assembled transcripts and known protein sequences from the SwissProt 2017_04 (The UniProt Consortium 2017) and Pfam 31.0 (El-Gebali et al. 2019) databases. We kept only SwissProt entries with confidence scores 1, 2 or 3 and we used the Pfam-A curated section of Pfam. We searched for sequence similarity using the blastx utility in the BLAST+2.8.1 package (Camacho et al. 2009; Altschul et al. 1990), keeping hits with maximum e-value 1e-3 and minimum protein sequence identity 40%, on repeat-masked cDNA sequences. We considered that transcripts were protein-coding if they overlapped with blastx hits over at least 150 bp. Genes were said to be protein-coding if at least one of their isoforms was classified as protein-coding, based on either the CSF score or on sequence similarity with known proteins.

### Long non-coding RNA selection

To construct a reliable lncRNA dataset, we selected newly-annotated genes classified as non-coding based on both the CSF score and on sequence similarity with known proteins and protein domains, as well as Ensembl-annotated genes with non-coding biotypes (“lincRNA”, “processed_transcript”, “antisense”, “TEC”, “macro_lncRNA”, “bidirectional_promoter_lncRNA”, “sense_intronic”). For newly detected genes, we applied several additional filters: we required a minimum exonic length (corresponding to the union of all annotated isoforms) of at least 200 bp for multi-exonic loci and of at least 500 bp for mono-exonic loci; we eliminated genes that overlapped for more than 5% of their exonic length with unmappable regions; we kept only loci that were classified as intergenic and at least 5 kb away from Ensembl-annotated protein-coding genes on the same strand; for multi-exonic loci, we required that all splice junctions be supported by reads with correct strand assignment (cf. above). For both *de novo* and Ensembl annotations, we removed transcribed loci that overlapped on at least 50% of their length with retrotransposed gene copies, annotated by the UCSC Genome Browser and from a previous publication (Carelli et al. 2016); we discarded loci that overlapped with UCSC-annotated tRNA genes and with RNA-type elements from RepeatMasker (Smit et al. 2003) on at least 25% of their length. We kept loci supported by at least 10 uniquely mapped RNA-seq reads and for which a ratio of sense to antisense transcription of at least 1% was observed in at least one sample. Although the fraction of reads stemming from the complementary mRNA strand, due to errors in library preparations, is very low in our samples (Supplementary Table 1), we noticed that loci situated on the antisense strand of highly expressed genes can have unreliable expression estimates, due to wrong strand-assignments of RNA-seq reads. Thus, for loci that had sense/antisense exonic overlap with other genes, we computed expression levels either on complete gene annotations, or only on exonic regions that had no overlap with other genes, and computed Spearman’s correlation coefficient between the two expression estimates, across all samples. We discarded loci for which the correlation coefficient was below 0.9.

### LncRNA and protein-coding gene promoter types

We assessed the proximity between the transcription start sites (TSS) of lncRNA loci and other genes TSS, to determine whether lncRNA promoters are unidirectional or bidirectional. We said that lncRNAs or protein-coding genes have bidirectional promoters if at least one of their annotated TSS is found within 1 kb of a different gene TSS. We also analyzed the proximity between promoter regions and Encode-annotated enhancers, in the mouse (Shen et al. 2012). We combined all enhancer coordinates across all tissues, and converted their coordinates between mm9 and mm10 genome assemblies with liftOver.

### Gene expression estimation

We computed the number of uniquely mapping reads unambiguously attributed to each gene using the Rsubread package in R (Liao et al. 2019), discarding reads that overlapped with multiple genes. We also estimated read counts and TPM (transcript *per* million) values *per* gene using Kallisto (Bray et al. 2016). To approach absolute expression levels estimates, for better comparisons across samples, we further normalized TPM values using a scaling approach (Brawand et al. 2011). Briefly, we ranked the genes in each sample according to their TPM values, we computed the variance of the ranks across all samples for each gene, and we identified the 100 least-varying genes, found within the inter-quartile range (25%-75%) in terms of average expression levels across samples. We derived normalization coefficients for each sample such that the median of the 100 least-varying genes be identical across samples. We then used these coefficients to normalize TPM values for each sample. We excluded mitochondrial genes from expression estimations and analyses, as these genes are highly expressed and can be variable across samples.

### Differential expression analyses

We used the DESeq2 (Love et al. 2014)(Smedley et al. 2009)(74)(75) package release 1.22.2 in R release 3.5.0 (R Core Team 2018) to test for differential expression across developmental stages, separately for each organ and species. We analyzed both protein-coding genes and lncRNAs, selected according to the criteria described above. We first performed a global differential expression analysis, using the likelihood ratio test to contrast a model including an effect of the developmental stage against the null hypothesis of homogeneous expression level across all developmental stages. This analysis was performed on all annotated protein-coding and lncRNA genes for each species, as well as on 1-to-1 orthologous genes for mouse and rat. In addition, we down-sampled the numbers of reads assigned to protein-coding genes to obtain identical average numbers of reads for protein-coding genes and lncRNAs. We also contrasted consecutive developmental stages, for each species and organ. For each test, we also computed the expression fold change based on average TPM values for each developmental stage/organ combination. To characterize expression variations across developmental stages, for DE genes shared between mouse and rat, TPM values were averaged across replicates and normalized by dividing by the maximum, for each species. The resulting relative expression profiles were combined across species and clustered with the K-means algorithm.

### Expression specificity index

We used the previously proposed tissue specificity index (Liao et al. 2006) to measure gene expression specificity across organs and developmental stages, provided by the formula: tau = sum (1 – r_i_)/(n-1), where r_i_ represents the ratio between the expression level in sample i and the maximum expression level across samples, and n represents the total number of samples. We computed this index on normalized TPM values, averaged across all replicates for a given species / organ / developmental stage combination (Supplementary Dataset 3).

### Gene markers for organs and developmental stages

We extracted protein-coding genes that had high expression specificity indexes (tau >= 0.85) in both mouse and rat, and for which the organ/stage in which maximum expression is observed is the same for both species. We selected genes that had an expression level (TPM) above 2 in at least one sample, for each species. For this analysis, we combined the young and aged adult samples.

### Gene ontology enrichment

We extracted gene ontology annotations for mouse, rat and chicken from the Ensembl 94 database, using BioMart (Smedley et al. 2015). We evaluated the enrichment of gene ontology categories between a target gene set (e.g., genes that differentially expressed between two developmental stages, in a given organ) and a background gene set (e.g., all genes expressed in that organ), using the hypergeometric test in R. We performed the gene ontology enrichment for the “biological process” category, and set the false discovery rate (FDR) threshold at 0.1. For the developmental transcription factor analysis presented in Figure 2, we selected genes associated with at least one of the following categories: multicellular organism development, system development, embryonic organ development, animal organ development or pattern specification process. We further selected from the resulting list those genes associated with at least one of the following categories: regulation of gene expression, gene expression, positive regulation of gene expression, negative regulation of gene expression, regulation of transcription by RNA polymerase II, regulation of transcription, DNA-templated.

### Homologous lncRNA family prediction

We used existing whole-genome alignments as a guide to predict homologous lncRNAs across species, as previously proposed (Washietl et al. 2014). We first constructed for each gene the union of its exon coordinates across all isoforms, hereafter termed “exon blocks”. We projected exon block coordinates between pairs of species using the liftOver utility and whole-genome alignments generated with blastz (http://www.bx.psu.edu/miller_lab/), available through the UCSC Genome Browser (Casper et al. 2018). To increase detection sensitivity, for the initial liftOver projection we required only that 10% of the reference bases remap on the target genome. Projections were then filtered, retaining only cases where the size ratio between the projected and the reference region was between 0.33 and 3 for mouse and rat (0.2 and 5 for comparisons involving chicken). To exclude recent lineage-specific duplications, regions with ambiguous or split liftOver projections were discarded. For genes where multiple exon blocks could be projected across species, we defined the consensus chromosome and strand in the target genome and discarded projected exon blocks that did not match this consensus. We then evaluated the order of the projected exon blocks on the target genes, to identify potential internal rearrangements. If internal rearrangements were due to the position of a single projected exon block, the conflicting exon block was discarded; otherwise, the entire projected gene was eliminated. As the projected reference gene coordinates could overlap with multiple genes in the target genome, we constructed gene clusters based on the overlap between projected exon block coordinates and target annotations, using a single-link clustering approach. We then realigned entire genomic loci for each pair of reference-target genes found within a cluster, using lastz (http://www.bx.psu.edu/miller_lab/) and the threaded blockset aligner (Blanchette et al. 2004). Using this alignment, we computed the percentage of exonic sequences aligned without gaps and the percentage of identical exonic sequence, for each pair of reference-target genes. We then extracted the best hit in the target genome for each gene in the reference genome based on the percentage of identical exonic sequence, requiring that the ratio between the maximum percent identity and the percent identity of the second-best hit be above 1.1. Reciprocal best hits were considered to be 1-to-1 orthologous loci between pairs of species. For analyses across all three species, we constructed clusters of reciprocal best hits from pairwise species comparisons, using a single-link clustering approach. Resulting clusters with more than 1 representative *per* species were discarded. To examine the validity of our procedure for homologous gene family prediction, we compared the resulting gene families with predictions from the Ensembl Compara pipeline (Herrero et al. 2016), extracted from Ensembl release 94, for protein-coding genes.

### Sequence evolution

We evaluated long-term evolutionary sequence conservation based on PhastCons (Siepel et al. 2005) scores, computed for the mouse genome using either a placental mammal or a vertebrate multiple species alignment available from the UCSC Genome Browser (Casper et al. 2018). We attributed a value of 0 to all genomic positions that are not present in the whole-genome alignment used for PhastCons computations. We computed average PhastCons scores on exonic sequences (excluding exonic regions overlapping with other genes), promoter regions (defined as 1 kb immediately upstream of the transcription start site, masking any exonic regions that overlapped in these regions) and splice sites (defined as the first two and last two bases of each intron). For genes with multiple promoters, we computed the average score across all promoters.

### Gene expression evolution

We computed global and *per-*gene expression level conservation between mouse and rat, for 1-to-1 orthologous genes. We first measured gene expression conservation for protein-coding genes and lncRNAs as a class. For each organ/developmental stage, we computed the expression level correlation between mouse and rat average TPM levels, across all orthologous pairs. We also computed the correlation between individuals within the same species; for organ/stages with more than two biological replicates we computed the average correlation coefficient across all possible pairs of individuals. We then evaluated the global extent of gene expression conservation through the ratio of the between-species correlation coefficient to the average within-species correlation coefficient.

Spearman’s rank correlation coefficients were used in all cases. We obtained confidence intervals for expression conservation measures through a bootstrap procedure, resampling 100 times the same number of genes with replacement. In addition to this global measure of expression conservation, we estimated the extent of between-species expression divergence *per* gene by computing Euclidean distances between relative expression profiles for each species. The relative expression profiles were derived from TPM values *per* organ/developmental stage, averaged across biological replicates, divided by the sum of all average TPM values.

### Statistical analyses and graphical representations

All statistical analyses and graphical representations were done with R (R Core Team 2018), version 3.5.0. We performed principal component analyses using the ade4 library (Dray and Dufour 2007) and hierarchical clustering of gene expression matrices using the hclust function in the stats package in R, on pairwise Euclidean distances. Principal component analyses were performed on log2-transformed expression levels, and expression profiles were centered and scaled prior to analysis. For all analyses involving multiple statistical tests, false discovery rates were computed with the Benjamini-Hochberg procedure (Benjamini and Hochberg 1995). 95% confidence intervals for median values of distributions were computed with the following formula: median +/-1.57 × IQR/sqrt(N), where IQR is the inter-quartile range, sqrt denotes the square root and N the number of points.

## Availability of data and materials

The raw and processed RNA-seq data were submitted to the NCBI Gene Expression Omnibus (GEO), under accession number GSE108348. Additional processed files and all scripts used to analyze the data are available at the address: ftp://pbil.univ-lyon1.fr/pub/datasets/Darbellay_LncEvoDevo.

## Author contributions

FD performed organ dissections, RNA extractions, quality control, prepared samples for sequencing and contributed to study design and manuscript preparation. AN designed the study, performed computational analyses and wrote the manuscript. All authors read and approved the final manuscript.

## Acknowledgements

We thank Denis Duboule and all members of the laboratory for advice and support, Mylène Docquier, Brice Petit and the Genomics Platform of the University of Geneva for RNA-seq data production, Amanda Cooksey and the Chickspress Team at the University of Arizona (http://geneatlas.arl.arizona.edu) for granting us access to chicken RNA-seq data, Ioannis Xenarios and the Vital-IT team for computational support. We thank Jean-Marc Matter (University of Geneva) for providing chicken eggs. This work was performed using the computing facilities of the CC LBBE/PRABI, the Vital-IT Center for high-performance computing of the SIB Swiss Institute of Bioinformatics (http://www.vital-it.ch) and the computing center of the French National Institute of Nuclear and Particle Physics (CC-IN2P3). This project was funded by the Swiss National Science Foundation (SNSF Ambizione grant PZ00P3_142636), the Agence Nationale pour la Recherche (ANR JCJC 2017 LncEvoSys). FD was supported by a FP7 IDEAL grant (259679).

---

**Supplementary Figure 1.**
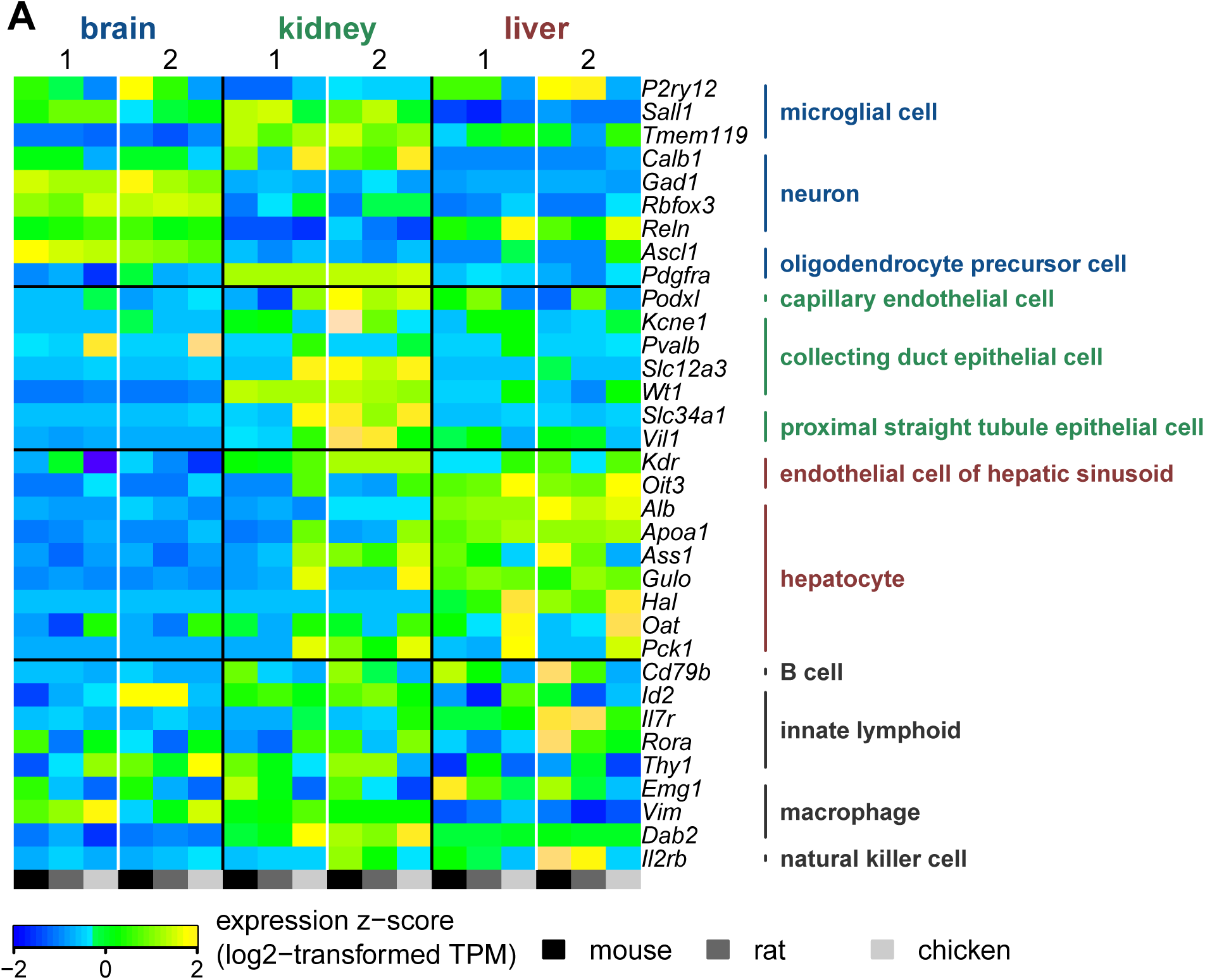

**Supplementary Figure 2.**
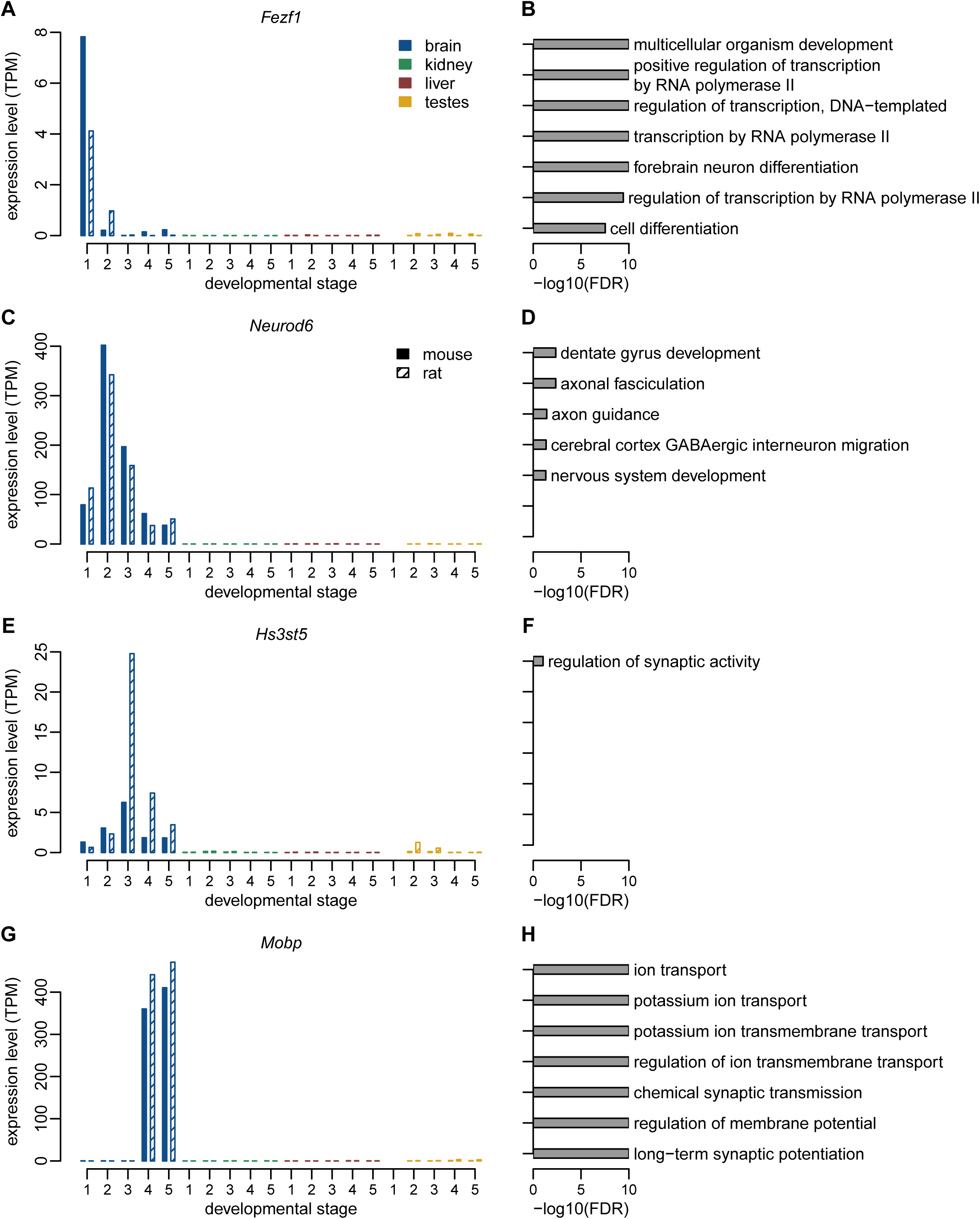

**Supplementary Figure 3.**
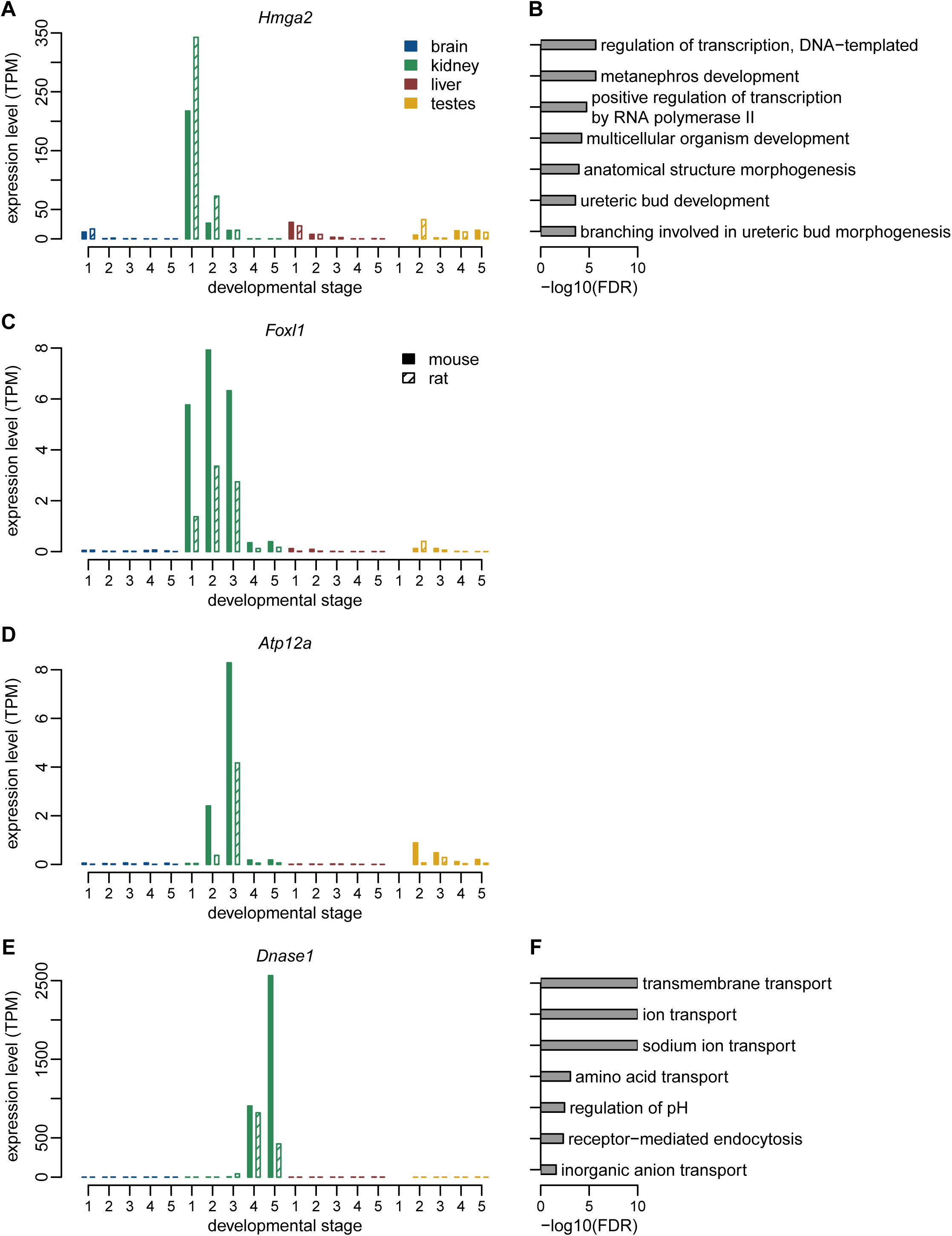

**Supplementary Figure 4.**
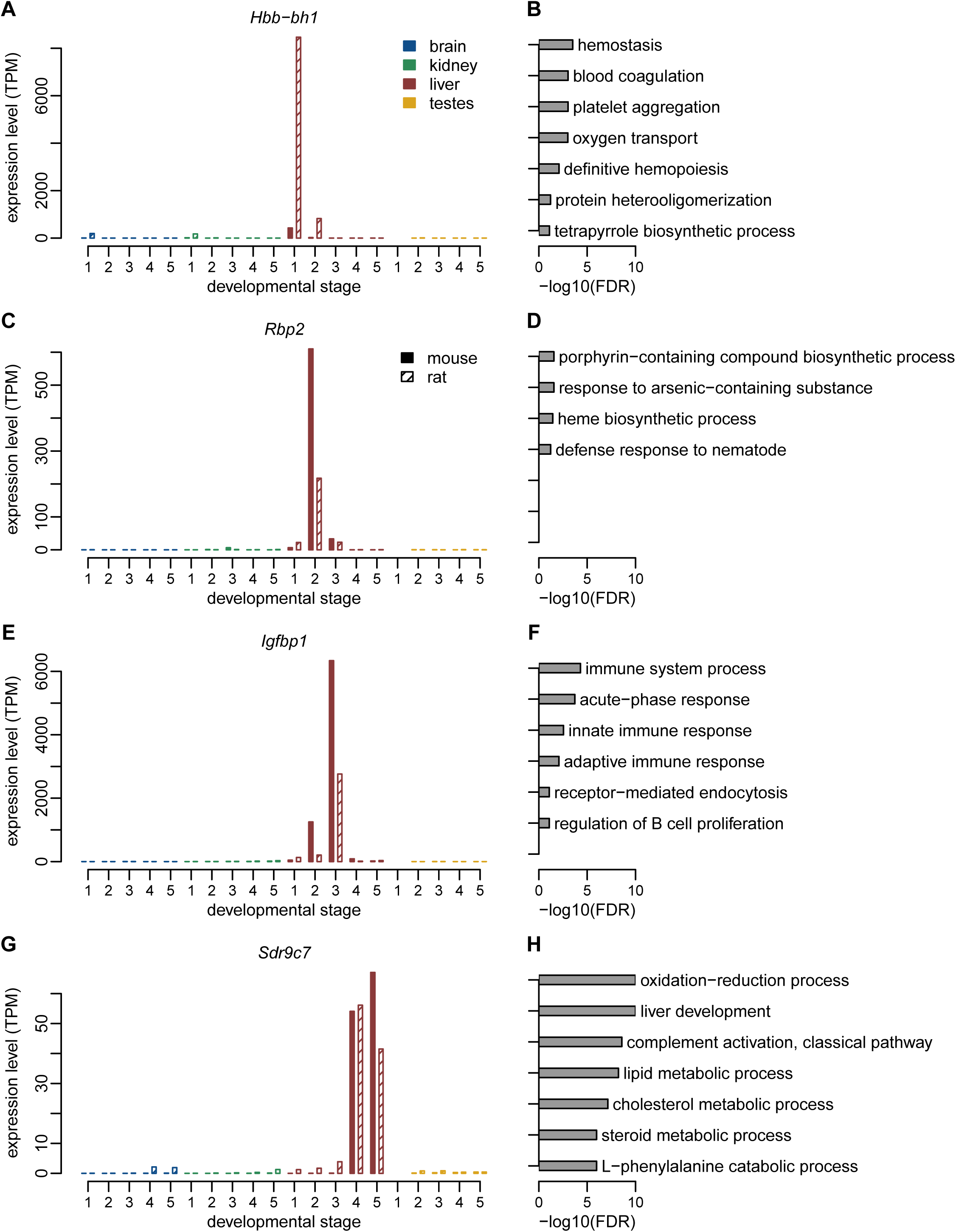

**Supplementary Figure 5.**
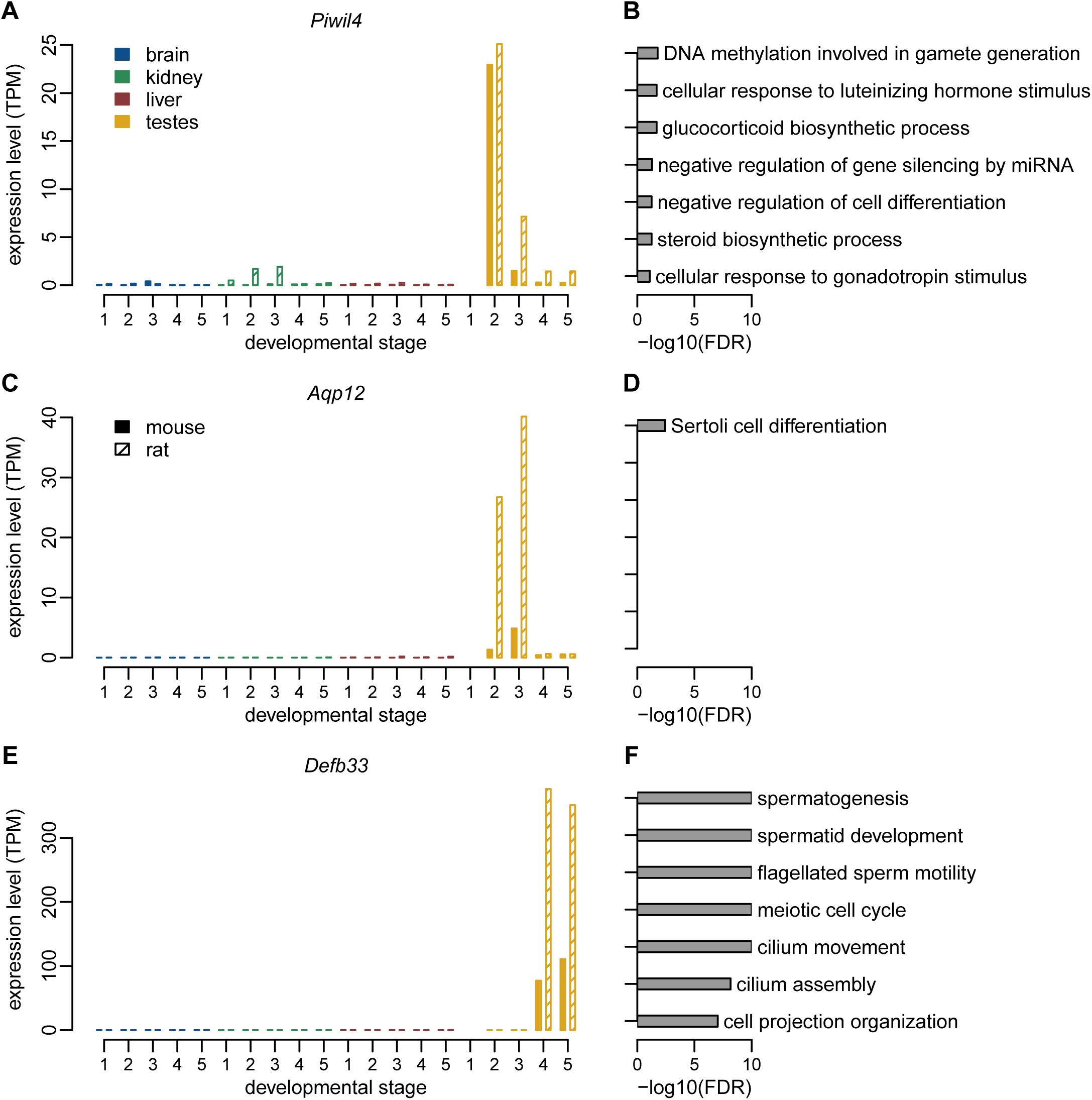

**Supplementary Figure 6.**
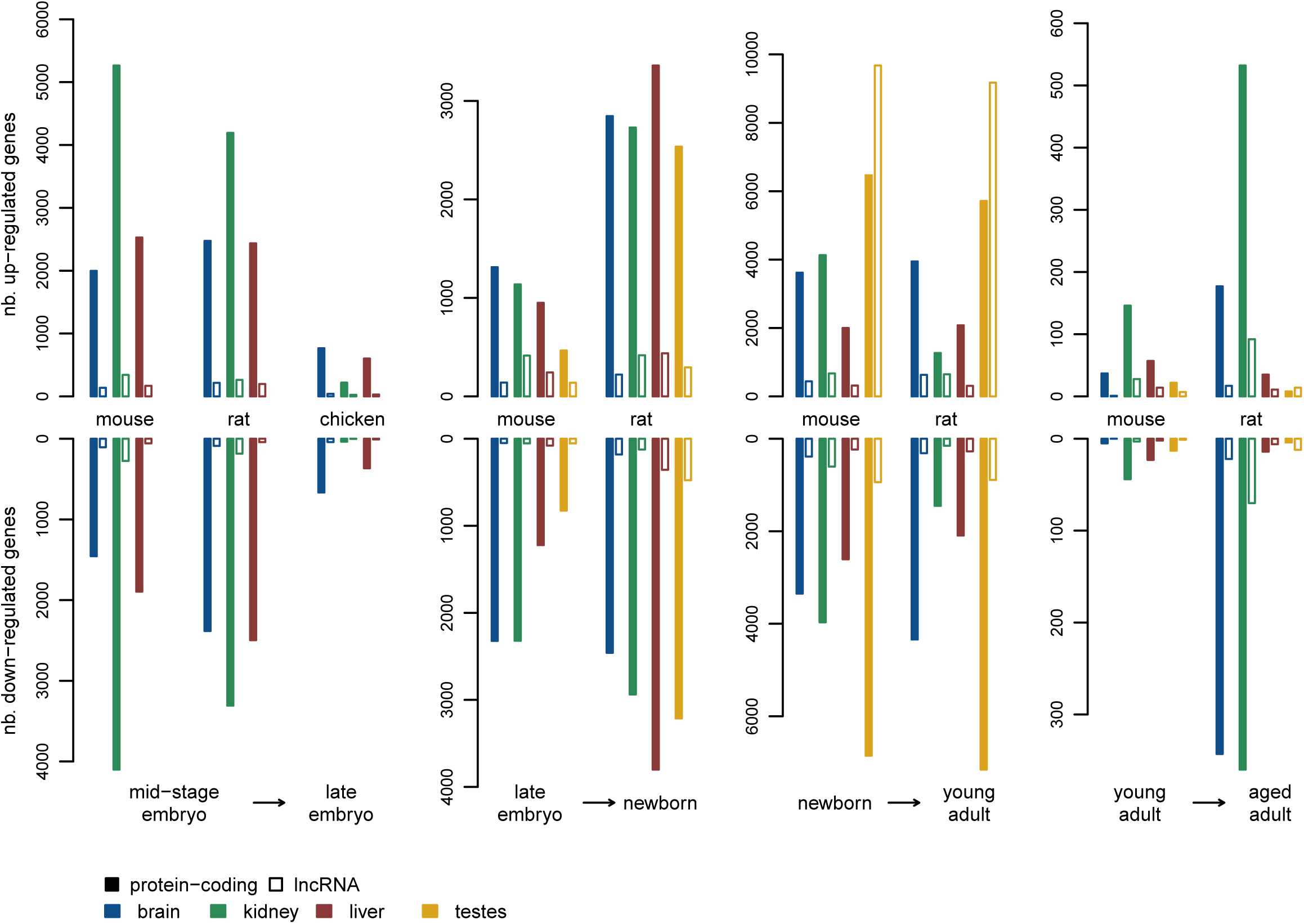

**Supplementary Figure 7.**
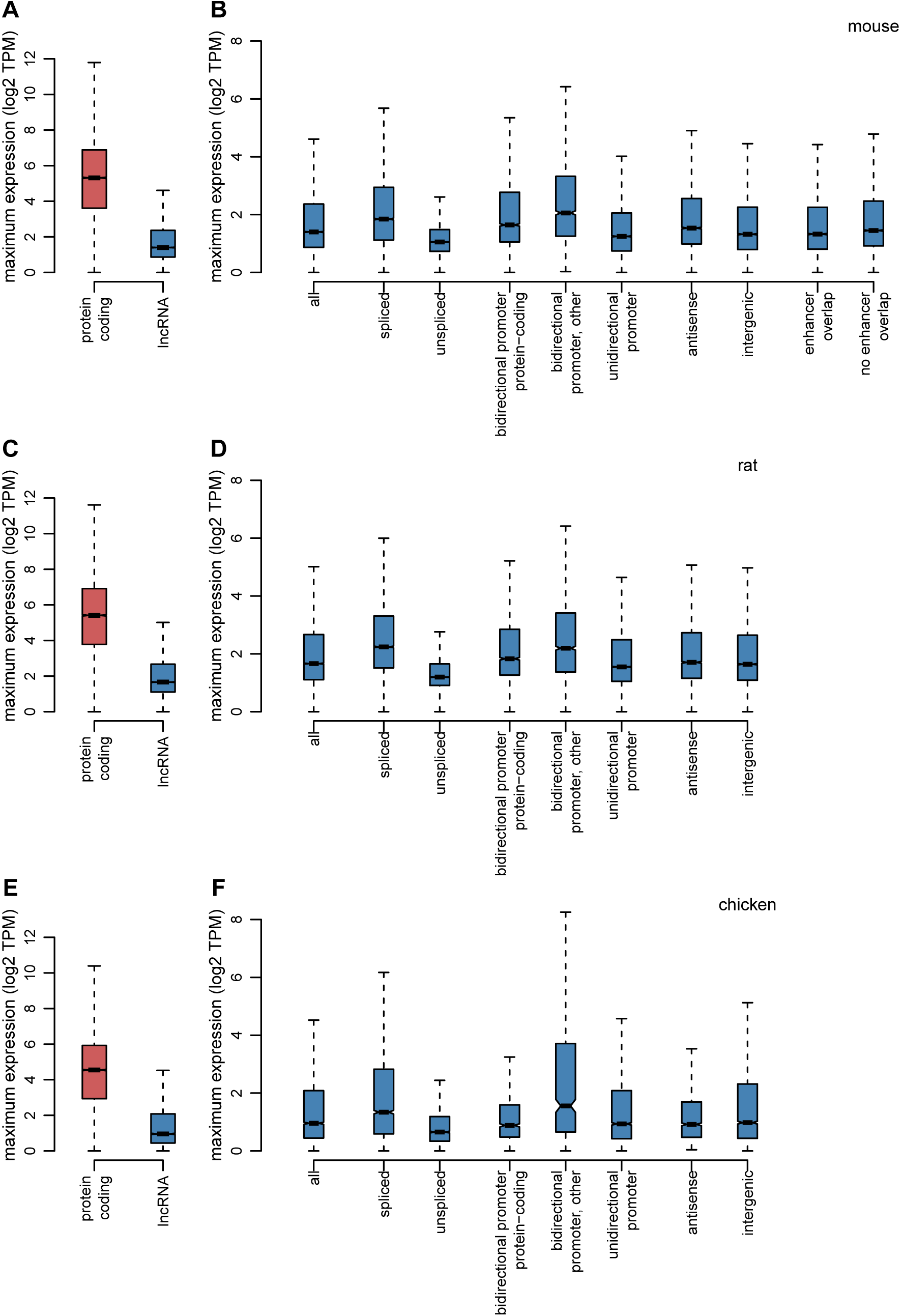

**Supplementary Figure 8.**
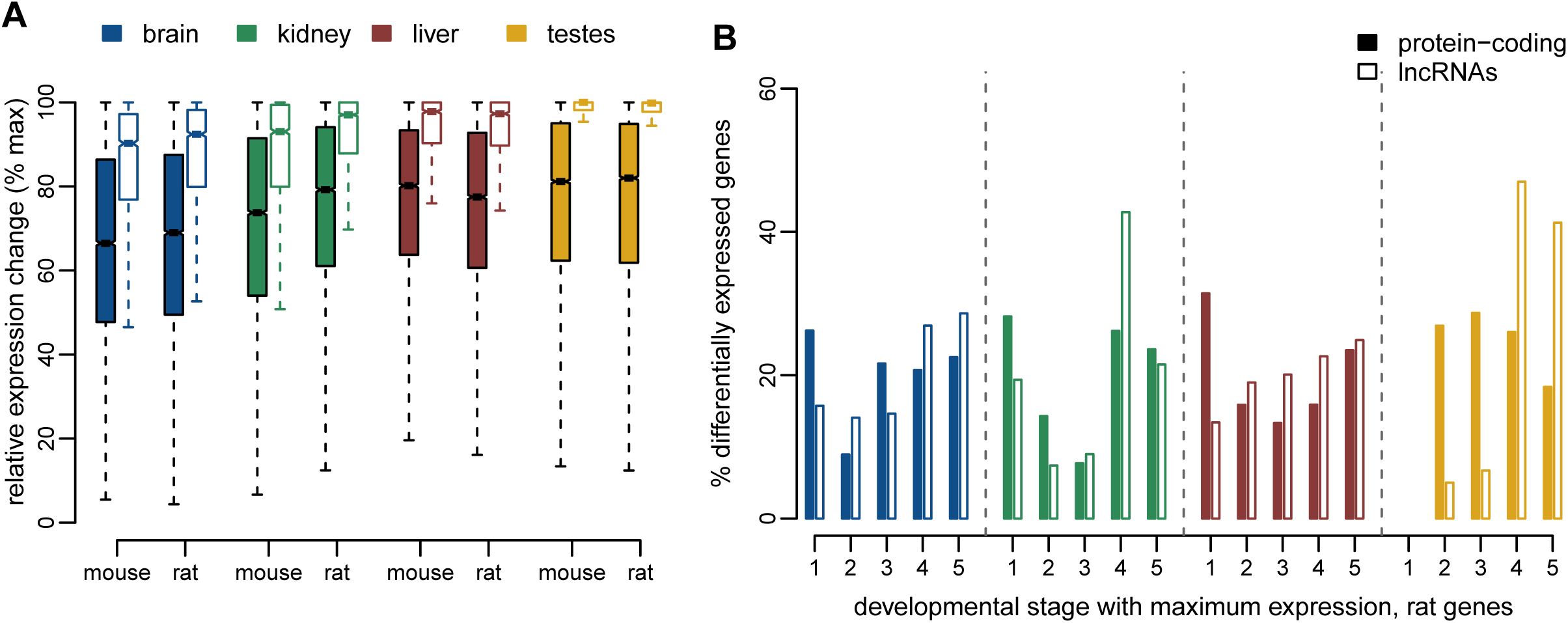

**Supplementary Figure 9.**
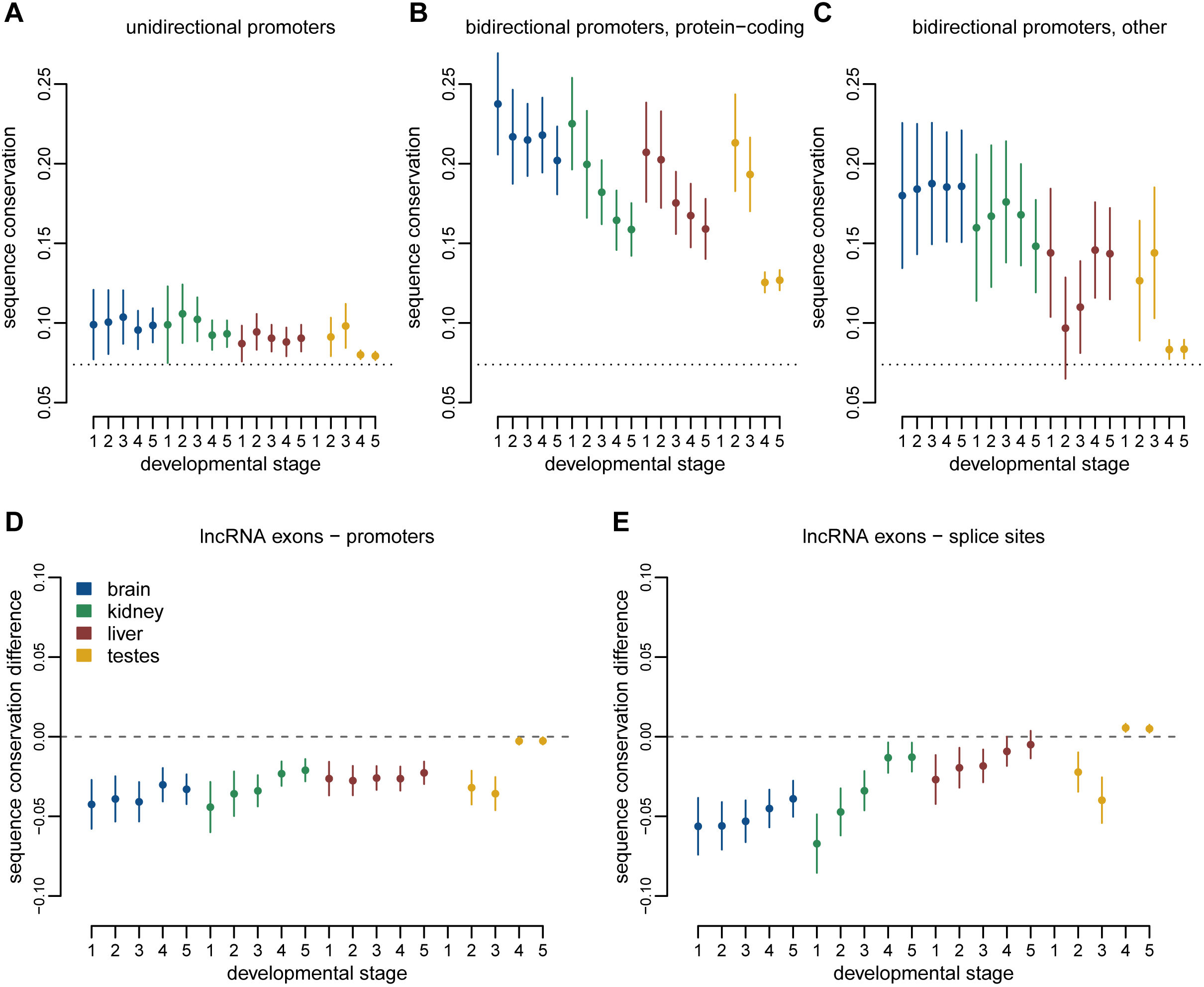

**Supplementary Figure 10.**
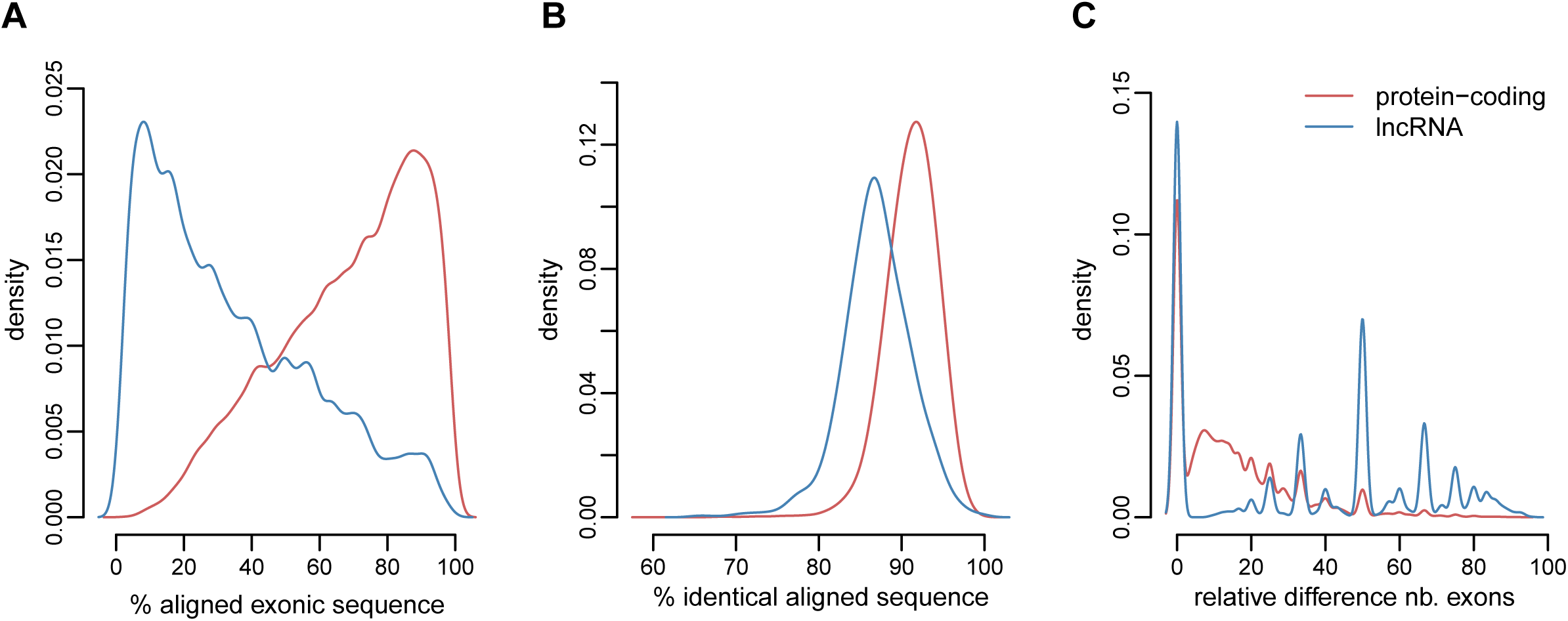

**Supplementary Figure 11.**
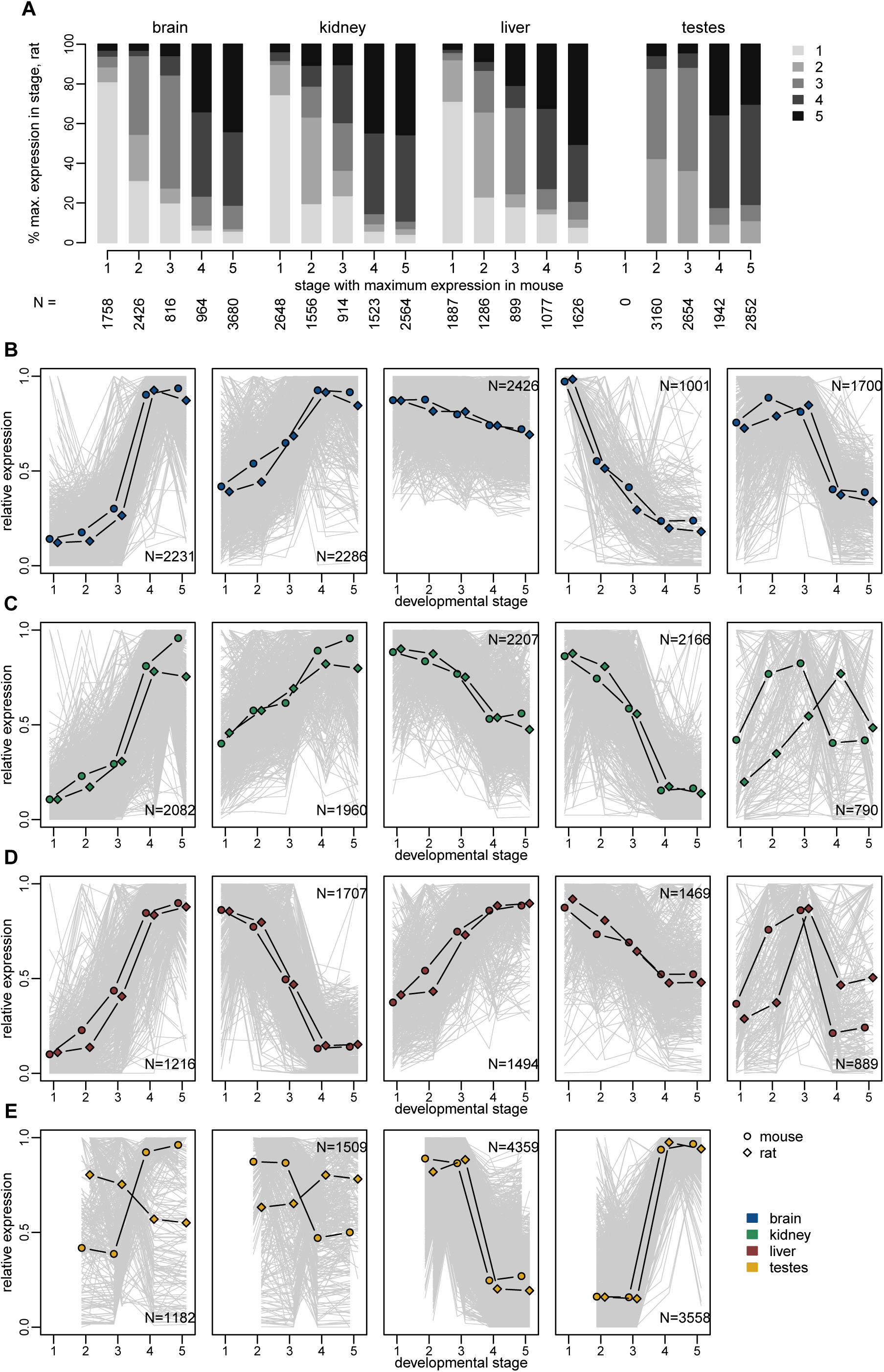

**Supplementary Figure 12.**
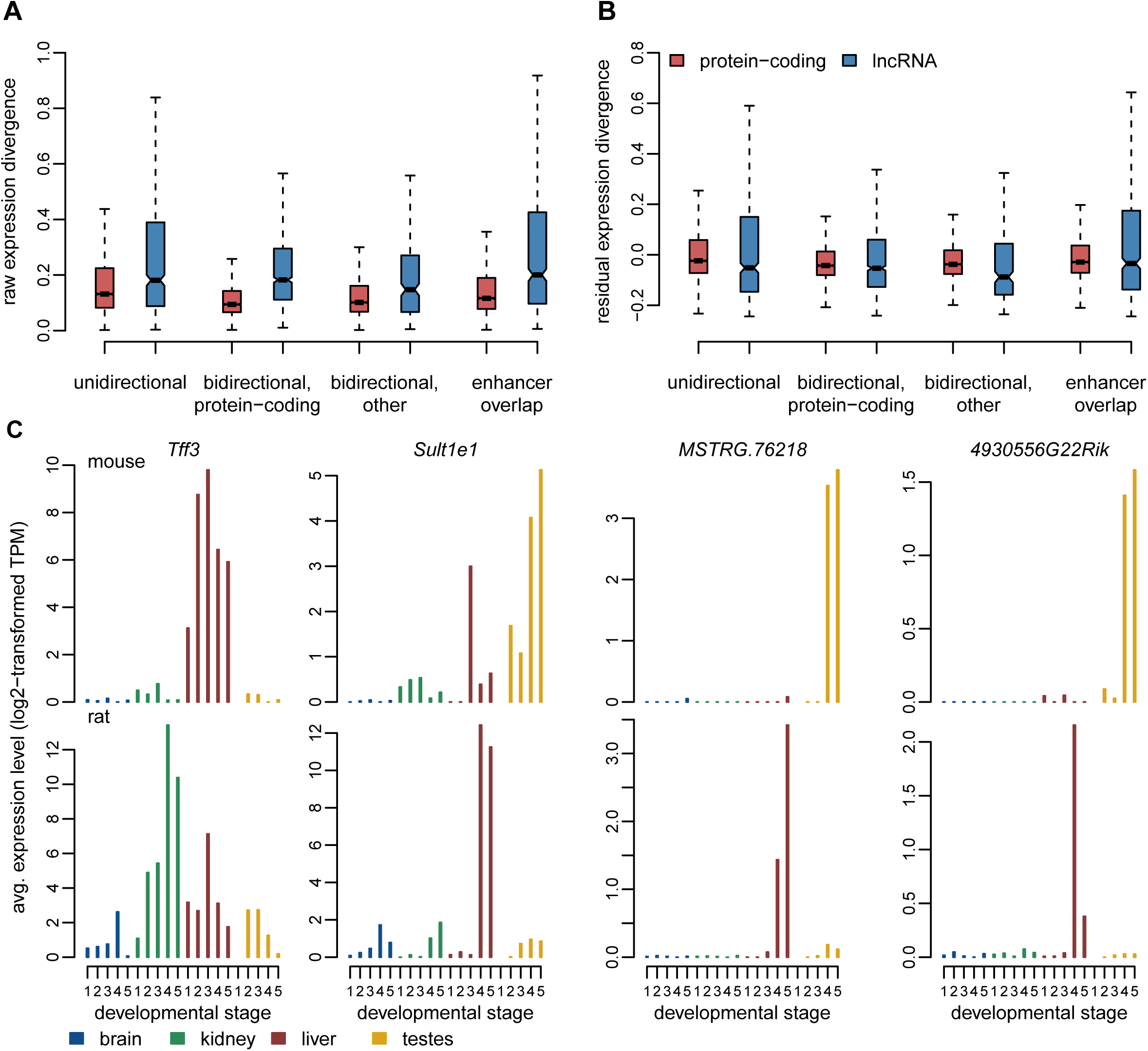

**Supplementary Figure 13.**
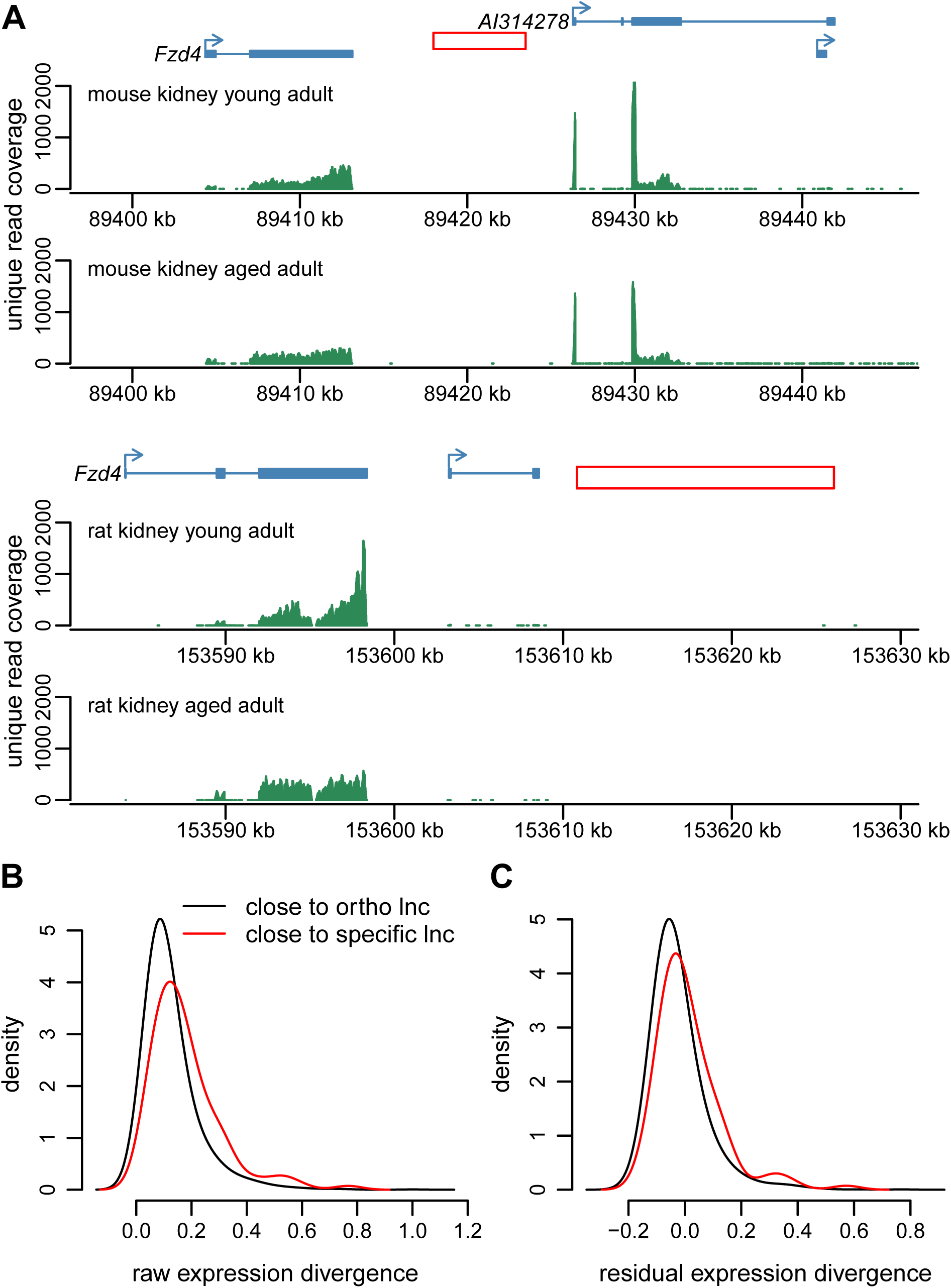

**Supplementary Figure 14.**
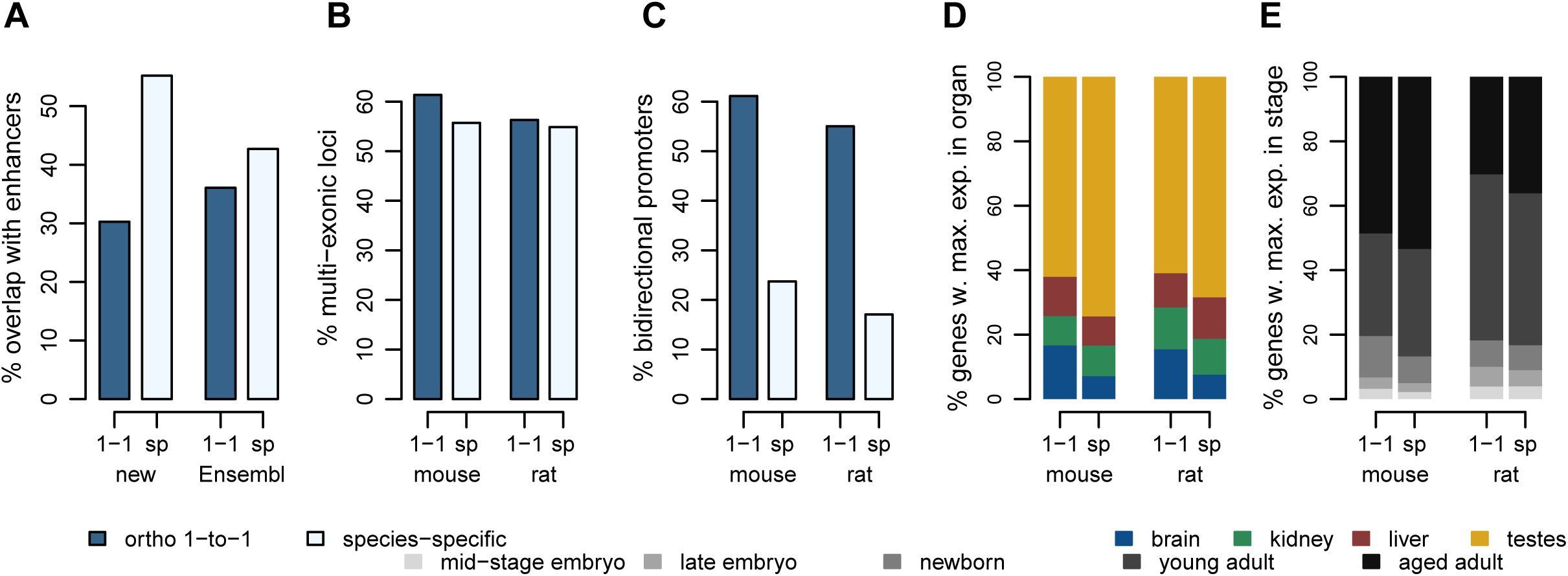

## References

Altschul SF, Gish W, Miller W, Myers EW, Lipman DJ. 1990. Basic local alignment search tool. J Mol Biol 215: 403–410.

Amândio AR, Necsulea A, Joye E, Mascrez B, Duboule D. 2016. Hotair is dispensible for mouse development. PLoS Genet 12: e1006232.

Anders S, Huber W. 2010. Differential expression analysis for sequence count data. Genome Biol 11: R106.

Anderson KM, Anderson DM, McAnally JR, Shelton JM, Bassel-Duby R, Olson EN. 2016. Transcription of the non-coding RNA upperhand controls Hand2 expression and heart development. Nature 539: 433–436.

Arendt D, Musser JM, Baker CVH, Bergman A, Cepko C, Erwin DH, Pavlicev M, Schlosser G, Widder S, Laubichler MD, et al. 2016. The origin and evolution of cell types. Nat Rev Genet 17: 744–757.

Ayers KL, Davidson NM, Demiyah D, Roeszler KN, Grützner F, Sinclair AH, Oshlack A, Smith CA. 2013. RNA sequencing reveals sexually dimorphic gene expression before gonadal differentiation in chicken and allows comprehensive annotation of the W-chromosome. Genome Biol 14.

Bassett AR, Akhtar A, Barlow DP, Bird AP, Brockdorff N, Duboule D, Ephrussi A, Ferguson-Smith AC, Gingeras TR, Haerty W, et al. 2014. Considerations when investigating lncRNA function in vivo. eLife 3: e03058.

Benjamini Y, Hochberg Y. 1995. Controlling the false discovery rate: a practical and powerful approach to multiple testing. J Roy Stat Soc B 57: 289+300.

Ben-Tabou de-Leon S, Davidson EH. 2007. Gene regulation: gene control network in development. Annu Rev Biophys Biomol Struct 36: 191.

Blanchette M, Kent WJ, Riemer C, Elnitski L, Smit AFA, Roskin KM, Baertsch R, Rosenbloom K, Clawson H, Green ED, et al. 2004. Aligning multiple genomic sequences with the threaded blockset aligner. Genome Res 14: 708–715.

Brannan CI, Dees EC, Ingram RS, Tilghman SM. 1990. The product of the H19 gene may function as an RNA. Mol Cell Biol 10: 28–36.

Brawand D, Soumillon M, Necsulea A, Julien P, Csárdi G, Harrigan P, Weier M, Liechti A, Aximu-Petri A, Kircher M, et al. 2011. The evolution of gene expression levels in mammalian organs. Nature 478: 343–348.

Bray NL, Pimentel H, Melsted P, Pachter L. 2016. Near-optimal probabilistic RNA-seq quantification. Nat Biotechnol 34: 525–527.

Brown CJ, Ballabio A, Rupert JL, Lafreniere RG, Grompe M, Tonlorenzi R, Willard HF. 1991. A gene from the region of the human X inactivation centre is expressed exclusively from the inactive X chromosome. Nature 349: 38–44.

Camacho C, Coulouris G, Avagyan V, Ma N, Papadopoulos J, Bealer K, Madden TL. 2009. BLAST+: architecture and applications. BMC Bioinformatics 10: 421.

Carelli FN, Hayakawa T, Go Y, Imai H, Warnefors M, Kaessmann H. 2016. The life history of retrocopies illuminates the evolution of new mammalian genes. Genome Res 26: 301–314.

Casper J, Zweig AS, Villarreal C, Tyner C, Speir ML, Rosenbloom KR, Raney BJ, Lee CM, Lee BT, Karolchik D, et al. 2018. The UCSC Genome Browser database: 2018 update. Nucleic Acids Res 46: D762–D769.

Cesana M, Cacchiarelli D, Legnini I, Santini T, Sthandier O, Chinappi M, Tramontano A, Bozzoni I. 2011. A long noncoding RNA controls muscle differentiation by functioning as a competing endogenous RNA. Cell 147: 358–369.

Cortez D, Marin R, Toledo-Flores D, Froidevaux L, Liechti A, Waters PD, Grützner F, Kaessmann H. 2014. Origins and functional evolution of Y chromosomes across mammals. Nature 508: 488–93.

Cunningham F, Achuthan P, Akanni W, Allen J, Amode MR, Armean IM, Bennett R, Bhai J, Billis K, Boddu S, et al. 2019. Ensembl 2019. Nucleic Acids Res 47: D745–D751.

Doolittle WF. 2018. We simply cannot go on being so vague about “function.” Genome Biol 19: 223.

Dray S, Dufour AB. 2007. The ade4 package: implementing the duality diagram for ecologists. J Stat Softw 22: 1–20.

El-Gebali S, Mistry J, Bateman A, Eddy SR, Luciani A, Potter SC, Qureshi M, Richardson LJ, Salazar GA, Smart A, et al. 2019. The Pfam protein families database in 2019. Nucleic Acids Res 47: D427–D432.

Engreitz JM, Haines JE, Perez EM, Munson G, Chen J, Kane M, McDonel PE, Guttman M, Lander ES. 2016. Local regulation of gene expression by lncRNA promoters, transcription and splicing. Nature 539: 452–455.

Gendrel A-V, Heard E. 2014. Noncoding RNAs and epigenetic mechanisms during X-chromosome inactivation. Annu Rev Cell Dev Biol 30: 561–580.

Goudarzi M, Berg K, Pieper LM, Schier AF. 2019. Individual long non-coding RNAs have no overt functions in zebrafish embryogenesis, viability and fertility. eLife 8.

Graur D, Zheng Y, Price N, Azevedo RBR, Zufall RA, Elhaik E. 2013. On the immortality of television sets: “function” in the human genome according to the evolution-free gospel of ENCODE. Genome Biol Evol 5: 578–590.

Green CD, Ma Q, Manske GL, Shami AN, Zheng X, Marini S, Moritz L, Sultan C, Gurczynski SJ, Moore BB, et al. 2018. A comprehensive roadmap of murine spermatogenesis defined by single-cell RNA-Seq. Dev Cell 46: 651–667.e10.

Groff AF, Sanchez-Gomez DB, Soruco MML, Gerhardinger C, Barutcu AR, Li E, Elcavage L, Plana O, Sanchez LV, Lee JC, et al. 2016. In vivo characterization of Linc-p21 reveals functional cis-regulatory DNA elements. Cell Rep 16: 2178–2186.

Grote P, Herrmann BG. 2015. Long noncoding RNAs in organogenesis: making the difference. Trends Genet TIG 31: 329–335.

Grote P, Wittler L, Hendrix D, Koch F, Währisch S, Beisaw A, Macura K, Bläss G, Kellis M, Werber M, et al. 2013. The tissue-specific lncRNA Fendrr is an essential regulator of heart and body wall development in the mouse. Dev Cell 24: 206–214.

Guttman M, Amit I, Garber M, French C, Lin MF, Feldser D, Huarte M, Zuk O, Carey BW, Cassady JP, et al. 2009. Chromatin signature reveals over a thousand highly conserved large non-coding RNAs in mammals. Nature 458: 223–227.

Haerty W, Ponting CP. 2013. Mutations within lncRNAs are effectively selected against in fruitfly but not in human. Genome Biol 14: R49.

Haerty W, Ponting CP. 2014. No gene in the genome makes sense except in the light of evolution. Annu Rev Genomics Hum Genet 15: 71–92.

Haerty W, Ponting CP. 2015. Unexpected selection to retain high GC content and splicing enhancers within exons of multiexonic lncRNA loci. RNA N Y N 21: 333–346.

Hamburger V, Hamilton HL. 1951. A series of normal stages in the development of the chick embryo. J Morphol 88: 49–92.

Herrero J, Muffato M, Beal K, Fitzgerald S, Gordon L, Pignatelli M, Vilella AJ, Searle SMJ, Amode R, Brent S, et al. 2016. Ensembl comparative genomics resources. Database J Biol Databases Curation 2016.

Hezroni H, Koppstein D, Schwartz MG, Avrutin A, Bartel DP, Ulitsky I. 2015. Principles of long noncoding RNA evolution derived from direct comparison of transcriptomes in 17 species. Cell Rep 11: 1110–1122.

Iyer MK, Niknafs YS, Malik R, Singhal U, Sahu A, Hosono Y, Barrette TR, Prensner JR, Evans JR, Zhao S, et al. 2015. The landscape of long noncoding RNAs in the human transcriptome. Nat Genet 47: 199–208.

Kaessmann H. 2010. Origins, evolution, and phenotypic impact of new genes. Genome Res 20: 1313–1326.

Kathleen Baxter K, Uittenbogaard M, Yoon J, Chiaramello A. 2009. The neurogenic basic helix-loop-helix transcription factor NeuroD6 concomitantly increases mitochondrial mass and regulates cytoskeletal organization in the early stages of neuronal differentiation. ASN Neuro 1.

Khalil AM, Guttman M, Huarte M, Garber M, Raj A, Rivea Morales D, Thomas K, Presser A, Bernstein BE, van Oudenaarden A, et al. 2009. Many human large intergenic noncoding RNAs associate with chromatin-modifying complexes and affect gene expression. Proc Natl Acad Sci U S A 106: 11667–11672.

Kim D, Langmead B, Salzberg SL. 2015. HISAT: a fast spliced aligner with low memory requirements. Nat Methods 12: 357–60.

Kutter C, Watt S, Stefflova K, Wilson MD, Goncalves A, Ponting CP, Odom DT, Marques AC. 2012. Rapid turnover of long noncoding RNAs and the evolution of gene expression. PLoS Genet 8: e1002841.

Latos PA, Pauler FM, Koerner MV, Şenergin HB, Hudson QJ, Stocsits RR, Allhoff W, Stricker SH, Klement RM, Warczok KE, et al. 2012. Airn transcriptional overlap, but not its lncRNA products, induces imprinted Igf2r silencing. Science 338: 1469–1472.

Liao B-Y, Scott NM, Zhang J. 2006. Impacts of gene essentiality, expression pattern, and gene compactness on the evolutionary rate of mammalian proteins. Mol Biol Evol 23: 2072–2080.

Liao Y, Smyth GK, Shi W. 2019. The R package Rsubread is easier, faster, cheaper and better for alignment and quantification of RNA sequencing reads. Nucleic Acids Res.

Lin MF, Carlson JW, Crosby MA, Matthews BB, Yu C, Park S, Wan KH, Schroeder AJ, Gramates LS, St Pierre SE, et al. 2007. Revisiting the protein-coding gene catalog of Drosophila melanogaster using 12 fly genomes. Genome Res 17: 1823–1836.

Lin MF, Jungreis I, Kellis M. 2011. PhyloCSF: a comparative genomics method to distinguish protein coding and non-coding regions. Bioinforma Oxf Engl 27: i275–282.

Liu SJ, Nowakowski TJ, Pollen AA, Lui JH, Horlbeck MA, Attenello FJ, He D, Weissman JS, Kriegstein AR, Diaz AA, et al. 2016. Single-cell analysis of long non-coding RNAs in the developing human neocortex. Genome Biol 17: 67.

Love MI, Huber W, Anders S. 2014. Moderated estimation of fold change and dispersion for RNA-seq data with DESeq2. Genome Biol 15: 550.

Marques AC, Hughes J, Graham B, Kowalczyk MS, Higgs DR, Ponting CP. 2013. Chromatin signatures at transcriptional start sites separate two equally populated yet distinct classes of intergenic long noncoding RNAs. Genome Biol 14: R131.

McMahon AP. 2016. Development of the Mammalian Kidney. Curr Top Dev Biol 117: 31–64.

Nakagaki BN, Mafra K, de Carvalho É, Lopes ME, Carvalho-Gontijo R, de Castro-Oliveira HM, Campolina-Silva GH, de Miranda CDM, Antunes MM, Silva ACC, et al. 2018. Immune and metabolic shifts during neonatal development reprogram liver identity and function. J Hepatol 69: 1294–1307.

Necsulea A, Soumillon M, Warnefors M, Liechti A, Daish T, Grutzner F, Kaessmann H. 2014. The evolution of lncRNA repertoires and expression patterns in tetrapods. Nature 505: 635–640.

Ørom UA, Derrien T, Beringer M, Gumireddy K, Gardini A, Bussotti G, Lai F, Zytnicki M, Notredame C, Huang Q, et al. 2010. Long noncoding RNAs with enhancer-like function in human cells. Cell 143: 46–58.

Pertea M, Pertea GM, Antonescu CM, Chang T-C, Mendell JT, Salzberg SL. 2015. StringTie enables improved reconstruction of a transcriptome from RNA-seq reads. Nat Biotechnol 33: 290–5.

Pertea M, Shumate A, Pertea G, Varabyou A, Breitwieser FP, Chang Y-C, Madugundu AK, Pandey A, Salzberg SL. 2018. CHESS: a new human gene catalog curated from thousands of large-scale RNA sequencing experiments reveals extensive transcriptional noise. Genome Biol 19: 208.

Ponjavic J, Ponting CP, Lunter G. 2007. Functionality or transcriptional noise? Evidence for selection within long noncoding RNAs. Genome Res 17: 556–65.

R Core Team. 2018. R: A Language and Environment for Statistical Computing. https://www.R-project.org/.

Ramsköld D, Wang ET, Burge CB, Sandberg R. 2009. An abundance of ubiquitously expressed genes revealed by tissue transcriptome sequence data. PLoS Comput Biol 5: e1000598.

Rinn JL, Kertesz M, Wang JK, Squazzo SL, Xu X, Brugmann SA, Goodnough LH, Helms JA, Farnham PJ, Segal E, et al. 2007. Functional demarcation of active and silent chromatin domains in human HOX loci by noncoding RNAs. Cell 129: 1311–1323.

Sauvageau M, Goff LA, Lodato S, Bonev B, Groff AF, Gerhardinger C, Sanchez-Gomez DB, Hacisuleyman E, Li E, Spence M, et al. 2013. Multiple knockout mouse models reveal lincRNAs are required for life and brain development. eLife 2: e01749.

Schüler A, Ghanbarian AT, Hurst LD. 2014. Purifying selection on splice-related motifs, not expression level nor RNA folding, explains nearly all constraint on human lincRNAs. Mol Biol Evol 31: 3164–3183.

Shen Y, Yue F, McCleary DF, Ye Z, Edsall L, Kuan S, Wagner U, Dixon J, Lee L, Lobanenkov VV, et al. 2012. A map of the cis-regulatory sequences in the mouse genome. Nature 488: 116–120.

Siepel A, Bejerano G, Pedersen JS, Hinrichs AS, Hou M, Rosenbloom K, Clawson H, Spieth J, Hillier LW, Richards S, et al. 2005. Evolutionarily conserved elements in vertebrate, insect, worm, and yeast genomes. Genome Res 15: 1034–50.

Smedley D, Haider S, Ballester B, Holland R, London D, Thorisson G, Kasprzyk A. 2009. BioMart--biological queries made easy. BMC Genomics 10: 22.

Smedley D, Haider S, Durinck S, Pandini L, Provero P, Allen J, Arnaiz O, Awedh MH, Baldock R, Barbiera G, et al. 2015. The BioMart community portal: an innovative alternative to large, centralized data repositories. Nucleic Acids Res 43: W589–598.

Smit AF., Hubley R, Green P. 2003. RepeatMasker Open-4.0. http://www.repeatmasker.org.

Soumillon M, Necsulea A, Weier M, Brawand D, Zhang X, Gu H, Barthès P, Kokkinaki M, Nef S, Gnirke A, et al. 2013. Cellular source and mechanisms of high transcriptome complexity in the mammalian testis. Cell Rep 3: 2179–2190.

Tabula Muris Consortium. 2018. Single-cell transcriptomics of 20 mouse organs creates a Tabula Muris. Nature 562: 367–372.

Theiler K. 1989. The house mouse: atlas of embryonic development. Springer-Verlag, Berlin Heidelberg https://www.springer.com/la/book/9783642884207.

The UniProt Consortium. 2017. UniProt: the universal protein knowledgebase. Nucleic Acids Res 45: D158–D169.

Uebbing S, Konzer A, Xu L, Backström N, Brunström B, Bergquist J, Ellegren H. 2015. Quantitative mass spectrometry reveals partial translational regulation for dosage compensation in chicken. Mol Biol Evol 32: 2716–2725.

Ulitsky I. 2016. Evolution to the rescue: using comparative genomics to understand long non-coding RNAs. Nat Rev Genet 17: 601–614.

Ulitsky I, Shkumatava A, Jan CH, Sive H, Bartel DP. 2011. Conserved function of lincRNAs in vertebrate embryonic development despite rapid sequence evolution. Cell 147: 1537–1550.

Villar D, Berthelot C, Aldridge S, Rayner TF, Lukk M, Pignatelli M, Park TJ, Deaville R, Erichsen JT, Jasinska AJ, et al. 2015. Enhancer evolution across 20 mammalian species. Cell 160: 554–566.

Washietl S, Kellis M, Garber M. 2014. Evolutionary dynamics and tissue specificity of human long noncoding RNAs in six mammals. Genome Res 24: 616–28.

Zakany J, Darbellay F, Mascrez B, Necsulea A, Duboule D. 2017. Control of growth and gut maturation by HoxD genes and the associated lncRNA Haglr. Proc Natl Acad Sci U S A 114: E9290–E9299.

