## Supplementary figure legends for "Comparative transcriptomics analyses across species, organs and developmental stages reveal functionally constrained lncRNAs"

*Supplementary Figure 1. Expression patterns of cell-type specific markers in mouse, rat and chicken samples.*

**A.** Expression of cell type-specific markers derived from single-cell experiments (full list provided in Supplementary Table 3), in our mouse, rat and chicken RNA-seq samples. The heatmap represents centered and scaled log2-transformed TPM levels (z-score). Developmental stages are indicated by numeric labels, 1 to 5. Average levels across biological replicates are shown. We show only organs and developmental stages that were sampled in all three species, for genes with 1-to-1 orthologues.

*Supplementary Figure 2. Genes with narrow expression distribution across organs and developmental stages, with maximum expression observed in brain.*

**A.** Expression pattern (TPM expression levels) across organs, species and developmental stages for *Fezf1*, which is predominantly expressed in brain mid-stage embryo, in both mouse and rat.

**B.** Gene ontology enrichment for genes with narrow expression distribution (organ/stage specificity index >= 0.85) and maximum expression reached in brain mid-stage embryo, in both mouse and rat.

**C.** Same as A, for *Neurod6*, which is predominantly expressed in brain late embryo.

**D.** Same as B, for genes with maximum expression in brain late embryo.

**E.** Same as A, for *Hs3st5*, which is predominantly expressed in brain newborn.

**F.** Same as B, for genes with maximum expression in brain newborn.

**G.** Same as A, for *Mobp*, which is predominantly expressed in brain young or aged adult.

**H.** Same as B, for genes with maximum expression in brain young or aged adult.

Full list of genes is provided in Supplementary Table 4.

*<u>Supplementary Figure 3. Genes with narrow expression distribution across organs and developmental stages, with maximum expression observed in kidney.</u>*

**A.** Expression pattern (TPM expression levels) across organs, species and developmental stages for *Hmga2*, which is predominantly expressed in kidney mid-stage embryo, in both mouse and rat.

**B.** Gene ontology enrichment for genes with narrow expression distribution (organ/stage specificity index >= 0.85) and maximum expression reached in kidney mid-stage embryo, in both mouse and rat.

**C.** Same as A, for *Foxl1*, which is predominantly expressed in kidney late embryo.

**D.** Same as A, for *Atp12a,* which is predominantly expressed in kidney newborn.

**E.** Same as A, for *Dnase1*, which is predominantly expressed in kidney young or aged adult.

**F.** Same as B, for genes with maximum expression in kidney young or aged adult.

Full list of genes is provided in Supplementary Table 4.

*<u>Supplementary Figure 4. Genes with narrow expression distribution across organs and developmental stages, with maximum expression observed in liver.</u>*

**A.** Expression pattern (TPM expression levels) across organs, species and developmental stages for *Hbb-bh1*, which is predominantly expressed in liver mid-stage embryo, in both mouse and rat.

**B.** Gene ontology enrichment for genes with narrow expression distribution (organ/stage specificity index >= 0.85) and maximum expression reached in liver mid-stage embryo, in both mouse and rat.

**C.** Same as A, for *Rbp2*, which is predominantly expressed in liver late embryo.

**D.** Same as B, for genes with maximum expression in liver late embryo.

**E.** Same as A, for *Igfbp1*, which is predominantly expressed in liver newborn.

**F.** Same as B, for genes with maximum expression in liver newborn.

**G.** Same as A, for *Sdr9c7*, which is predominantly expressed in liver young or aged adult.

**H.** Same as B, for genes with maximum expression in liver young or aged adult.

Full list of genes is provided in Supplementary Table 4.

*Supplementary Figure 5. Genes with narrow expression distribution across organs and developmental stages, with maximum expression observed in testes.*

**A.** Expression pattern (TPM expression levels) across organs, species and developmental stages for *Piwil4*, which is predominantly expressed in testes late embryo, in both mouse and rat.

**B.** Gene ontology enrichment for genes with narrow expression distribution (organ/stage specificity index >= 0.85) and maximum expression reached in testes late embryo, in both mouse and rat.

**C.** Same as A, for *Aqp12*, which is predominantly expressed in testes newborn.

**D.** Same as B, for genes with maximum expression in testes newborn.

**E.** Same as A, for *Defb33*, which is predominantly expressed in testes young or aged adult.

**F.** Same as B, for genes with maximum expression in testes young or aged adult.

Full list of genes is provided in Supplementary Table 4.

*Supplementary Figure 6. Number of differentially expressed genes between consecutive developmental stages.*

Numbers of significantly up-regulated and down-regulated protein-coding genes and lncRNAs (FDR<0.01). Differential expression tests are performed separately for each organ and species, comparing consecutive developmental stages.

*Supplementary 7. Expression level distribution for different classes of lncRNAs, for mouse, rat and chicken.*

**A.** Distribution of the maximum expression level (log2-transformed TPM), for mouse protein-coding genes and lncRNAs.

**B.** Distribution of the maximum expression level, for different classes of lncRNAs, in the mouse. From left to right: all lncRNAs, spliced (multi-exonic) lncRNAs, unspliced (mono-exonic) lncRNAs, lncRNAs with bidirectional promoters shared with protein-coding genes, lncRNAs with bidirectional promoters shared with other types of genes, lncRNA with unidirectional promoters, antisense lncRNAs (that have exonic or intronic overlap with protein-coding genes on the opposite strand), intergenic lncRNAs (that have no overlap with protein-coding genes on the opposite strand), lncRNAs that have an Encode-annotated enhancer within 1kb of their transcription start site, lncRNAs that are further away from Encode-annotated enhancers.

**C.** Same as A, for the rat.

**D.** Same as B, for the rat. For this species, we did not analyze the proximity between lncRNA promoters and enhancers, for lack of data.

**E.** Same as A, for the chicken.

**F.** Same as B, for the chicken. For this species, we did not analyze the proximity between lncRNA promoters and enhancers, for lack of data.

*Supplementary Figure 8. Patterns of differential expression among developmental stages for protein-coding genes and lncRNAs.*

**A.** Distribution of the relative expression change, defined as the difference between the maximum and the minimum expression level across developmental stages, normalized by the maximum expression level, for mouse and rat protein-coding genes and lncRNAs that are significantly differentially expressed (FDR<0.01) among developmental stages. Higher values indicate higher fold expression changes.

**B.** Distribution of the developmental stage in which the maximum expression is observed, for rat protein-coding genes and lncRNAs that are significantly differentially expressed (FDR<0.01) among developmental stages.

**A,B.** Differential expression analyses are performed separately for each organ.

*Supplementary Figure 9. Estimates of long-term sequence conservation scores for different regions of lncRNA loci.*

**A.** Distribution of the promoter sequence conservation score (PhastCons) for lncRNAs that have unidirectional promoters (no other transcription start site within 1kb of the start of the locus), and which are significantly expressed (TPM>=1) in each organ and developmental stage. Precomputed PhastCons score for placental mammals were provided by the UCSC Genome Browser. Exonic regions that overlap with other genes were masked. Dots represent median values, vertical bars represent 95% confidence intervals.

**B.** Same as A, for lncRNAs that have bidirectional promoters shared with protein-coding genes.

**C.** Same as A, for lncRNAs that have bidirectional promoters shared with other types of genes (non-coding).

**D.** Distribution of the difference between the exonic sequence conservation (PhastCons) score and the promoter score, for mouse lncRNAs that are significantly expressed (TPM>=1) in each organ and developmental stage.

**E.** Same as D, for the difference between exonic and splice site sequence conservation score.

*Supplementary Figure 10. Gene structure conservation for orthologous lncRNAs.*

**A.** Distribution of the percentage of exonic sequence aligned without gaps, with respect to the maximum exonic length, for each pair of orthologous genes, between mouse and rat. Red: protein-coding genes, blue: lncRNAs.

**B.** Distribution of the percentage of identical exonic sequence, with respect to the exonic sequence length aligned without gaps, for each pair of orthologous genes. Red: protein-coding genes, blue: lncRNAs.

**C.** Distribution of the relative difference of the number of exons, defined as the absolute difference of the number of exons of the two species, divided by the maximum number of exons across species, for each pair of orthologous genes. Red: protein-coding genes, blue: lncRNAs.

*Supplementary Figure 11. Conservation of developmental expression profiles between mouse and rat, for orthologous protein-coding genes.*

**A**. Comparison of the developmental stage in which maximum expression is observed, for orthologous protein-coding genes that are significantly differentially expressed (FDR<0.01) among developmental stages, for both mouse and rat. Genes are divided based on the developmental stage where maximum expression is observed in mouse organs (X-axis). The Y axis represents the percentage of orthologous genes that reach maximum expression in each developmental stage, in the rat. Numbers of analyzed genes are provided below the plot.

**B.** Expression profiles of orthologous protein-coding genes that are significantly differentially expressed (FDR<0.01) among developmental stages, for both mouse and rat, in the brain. TPM values were averaged across replicates and normalized by dividing by the maximum, for each species. The resulting relative expression profiles were combined across species and clustered with the K-means algorithm. The average profiles of the genes belonging to each cluster are shown. Gray lines represent profiles of individual genes from a cluster. Numbers of genes in each cluster are shown in the plot.

**C.** Same as B, for the kidney.

**D.** Same as B, for the liver.

**E.** Same as B, for the testes. For this organ, we searched for only 4 clusters with the K-means algorithm.

*Supplementary Figure 12. Expression pattern divergence between mouse and rat.*

**A.** Distribution of raw expression divergence values for different classes of protein-coding genes and lncRNAs, depending on their promoter type in mouse: unidirectional, bidirectional shared with protein-coding genes, bidirectional shared with non-coding genes, overlap with Encode-annotated enhancers.

**B.** Same as A, for the residual expression divergence values, after correction for the average expression levels.

**C.** Examples of expression profiles in mouse and rat, for the top 2 most-divergent protein-coding and lncRNA genes.

*Supplementary Figure 13. Candidate species-specific lncRNAs.*

**A.** Genomic localization and RNA-seq read coverage of a candidate mouse-specific lncRNA, situated downstream of the *Fzd4* gene. RNA-seq data is shown for young and aged adult kidney.

**B.** Distribution of the raw expression divergence for protein-coding genes that are transcribed from the same bidirectional promoters as lncRNAs with 1-to-1 orthologues in mouse and rat (black), or as candidate species-specific lncRNAs (red).

**C.** Same as A, after correcting the expression divergence for the average expression level.

*Supplementary Figure 14. Genomic and expression characteristics of candidate species-specific lncRNAs.*

**A.** Percentage of mouse lncRNAs for which the predicted transcription start site is found within 1kb of an Encode-annotated enhancer. LncRNAs are divided into loci with predicted 1-to-1 orthologues in the rat (1-1, dark blue) and mouse-specific lncRNAs (sp, light blue). LncRNAs are further separated into newly-annotated (new) or previously known (Ensembl).

**B.** Same as A, for the percentage of multi-exonic loci, for mouse and rat.

**C.** Same as A, for the percentage of loci that have predicted bidirectional promoters, for mouse and rat.

**D.** Distribution of the organ in which maximum expression is observed, for mouse and rat lncRNAs. LncRNAs are divided into loci with predicted 1-to-1 orthologues (1-1) and species-specific lncRNAs (sp).

**E.** Same as D, for the distribution of the developmental stage in which maximum expression is observed.

**Supplementary Table List**

**Supplementary Table 1.** List of RNA-seq samples generated specifically for this project, and used for all downstream expression analyses.

**Supplementary Table 2.** List of additional, previously published RNA-seq samples, included in the lncRNA detection pipeline.

**Supplementary Table 3.** Cell-type markers for the four organs analyzed here, derived from single-cell transcriptomics analyses.

**Supplementary Table 4.** List of putative organ/developmental stage markers. This list contains protein-coding genes with narrow expression distribution (organ/stage specificity value >= 0.85), which reach their maximum expression in the same organ and stage for mouse and rat.

**Supplementary Table 5.** Numbers of protein-coding genes and lncRNAs that have an average TPM expression level of at least 1 in each organ / developmental stage combination, for each species.

**Supplementary Table 6.** Sequence conservation scores (average PhastCons scores), for exons, introns, promoters and splice sites, for mouse protein-coding genes and lncRNAs.

**Supplementary Table 7.** List of 30 lncRNAs that are predicted to be 1-to-1 orthologues in mouse, rat and chicken.

**Supplementary Table 8.** Expression pattern and sequence conservation scores for protein-coding genes and lncRNAs, for mouse and rat 1-to-1 orthologues.

**Supplementary Dataset List**

README files are provided within each supplementary dataset archive, explaining the file lists.

**Supplementary Dataset 1.** Complete gene annotations for mouse, rat and chicken.

**Supplementary Dataset 2.** Gene expression levels (raw and normalized TPM values, unique read counts).

**Supplementary Dataset 3.** Expression patterns (average across replicates, samples with maximum expression), expression specificity indexes, lists of organ/developmental stage markers, gene ontology enrichment for organ/developmental stage markers.

**Supplementary Dataset 4.** Results of the differential expression analyses across all developmental stages, or between consecutive developmental stages, for each organ and each species.

**Supplementary Dataset 5.** Predicted orthologous gene families and sequence conservation statistics.

**Supplementary Dataset 6.** Raw and normalized expression values (TPM) for orthologous protein-coding and lncRNA families.

**Supplementary Dataset 7.** Expression pattern divergence for mouse and rat orthologous genes.

**Supplementary Dataset 8.** Lists of candidate species-specific lncRNAs.

**Supplementary figure legends**

**Supplementary table legends**
